# Replisomes restrict SMC-mediated DNA-loop extrusion *in vivo*

**DOI:** 10.1101/2025.02.23.639750

**Authors:** Qin Liao, Hugo B. Brandão, Zhongqing Ren, Xindan Wang

## Abstract

Structural maintenance of chromosomes (SMC) complexes organize genomes by extruding DNA loops, while replisomes duplicate entire chromosomes. These essential molecular machines must collide frequently in every cell cycle, yet how such collisions are resolved *in vivo* remains poorly understood. Taking advantage of the ability to load SMC complexes at defined sites in the *Bacillus subtilis* genome, we engineered head-on and head-to-tail collisions between SMC complexes and the replisome. Replisome progression was monitored by marker frequency analysis, and SMC translocation was monitored by time-resolved ChIP-seq and Hi-C. We found that SMC complexes do not impede replisome progression. By contrast, replisomes restrict SMC translocation regardless of collision orientations. Combining experimental data with simulations, we determined that SMC complexes are blocked by the replisome and then released from the chromosome. Occasionally, SMC complexes can bypass the replisome and continue translocating. Our findings establish that the replisome is a barrier to SMC-mediated DNA-loop extrusion *in vivo*, with implications for processes such as chromosome segregation, DNA repair, and gene regulation that require dynamic chromosome organization in all organisms.

## Introduction

Structural Maintenance of Chromosome (SMC) complexes play a key role in shaping the three-dimensional organization of genomes from bacteria to humans ^1, 2, 3^. The SMC complex binds to DNA, captures a small DNA loop, and enlarges the loop progressively by reeling the flanking DNA regions into the loop ^4, 5, 6, 7^. This process, known as DNA-loop extrusion, is a conserved mechanism for SMC actions. Through loop extrusion, SMC complexes generate interactions of DNA segments, which has been shown to contribute to many processes such as chromosome organization and segregation, gene expression, DNA replication, DNA recombination and repair ^8, 9^. In cells, SMC complexes are capable of traversing thousands to millions of DNA base pairs during loop extrusion, and will inevitably encounter other DNA-bound molecules on the crowded chromatin fiber.

One crucial machinery that the SMC complex encounters is the replisome, which duplicates entire chromosomes one nucleotide at a time. In eukaryotes, the SMC cohesin complexes load on the chromosomes during interphase and compact the DNA into topologically associating domains ^10, 11, 12^. During DNA replication, cohesin complexes establish sister chromatid cohesion, which is important for proper chromosome segregation ^13, 14, 15^. Thus, cohesins and replisomes collide frequently during replication ^16, 17, 18, 19^. However, the *in vivo* consequence of cohesin-replisome collisions and the effect of these collisions on DNA replication and chromosome folding are unknown. One major challenge to investigate these problems in eukaryotes *in vivo* is the lack of defined sites for replisome-cohesin collisions. This complexity motivated us to use a bacterial model, which has defined replication origin, defined SMC loading sites, and definable replisome-SMC collision sites, to dissect the fundamental principles governing the rules of engagement ^20^ between SMC and the replisome in cells.

In bacteria, DNA replication starts at a single origin and proceeds bidirectionally to the terminus region. Throughout the replication cycle, the SMC complex preferentially loads onto the *parS* sites by the partitioning protein ParB ^21, 22, 23, 24^. Once loaded, SMC travels processively from *parS* to the terminus and juxtaposes the left and right chromosomal arms ^25, 26^. This process connects DNA *in cis* and helps separate the newly replicated sister chromosomes ^27, 28^ (**Figure 1A**). Most bacterial genomes possess multiple *parS* sites and they all have varying distances from the replication origin ^29^. Because SMC and the replisome start their movement at different sites, collisions will occur frequently: SMC and replisome traveling towards each other will generate head-on collisions; SMC and replisome traveling in the same direction may generate head-to-tail collisions. Failures to resolve these collisions may have severe consequences on DNA replication and segregation. For instance, stalled or collapsed replication forks by SMC will perturb DNA replication, and premature unloading of SMC by the replisome will impair chromosome segregation. Therefore, SMC-replisome collisions have strong implications for genome integrity but the rules of engagement between SMC and the replisome in bacterial cells remain unclear.

**Figure 1.**
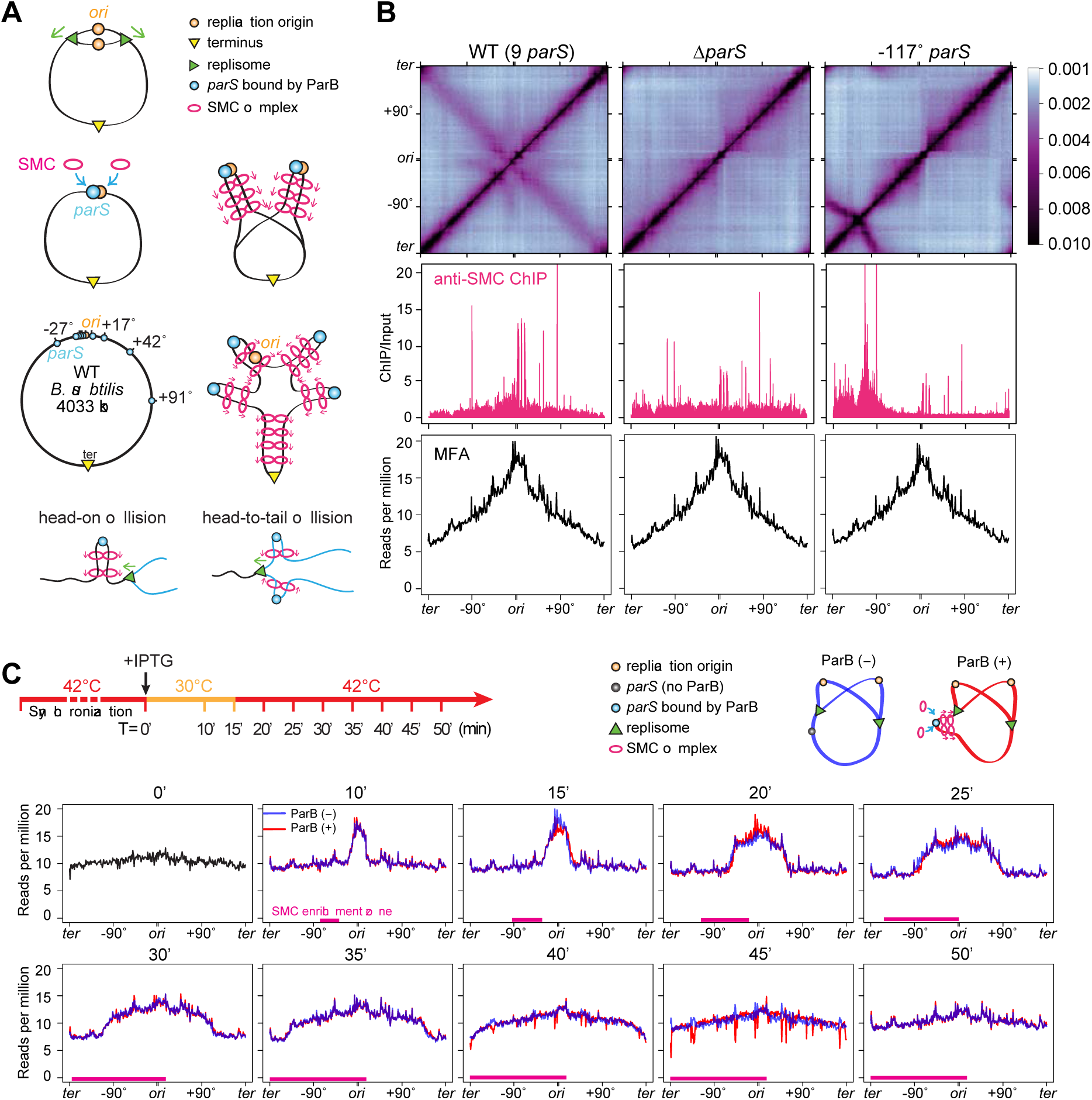
SMC translocation does not affect replication progression. See also Figures S1 and S2. **(A)** Schematics depicting replisome progression (top row), SMC loading and translocation helping origin segregation (2nd row), location of *parS* sites in WT *B. subtilis* and their effect on SMC loading (3rd row), and head-on and head-to-tail collisions between SMC and the replisome (bottom row). **(B)** Hi-C maps (top row), ChIP enrichments of SMC (middle row) and marker frequency analysis (MFA) plots (bottom row) of exponentially growing *B. subtilis* strains containing wild-type *parS* sites (PY79, left) ^32^, no *parS* site (BWX3212, middle) ^25^, or a single *parS* site at -117° (BWX3381, right) ^25^. **(C)** Time-course MFA plots of a *dnaB*(ts) strain containing a single *parS* site at -59° and IPTG-inducible copy of *parB* as the sole source of SMC loader protein (BWX5297). As indicated in the experimental timeline (top row), cells were arrested at G1 by growing at non-permissive temperature (42°C) for 45 min, then shifted to permissive temperature (30°C) for 15min, and finally back to 42°C for the rest of time to allow only one round of synchronized replication. See more analysis in **Figure S1**. Cells were grown with (red) or without (blue) IPTG at the onset of replication (T = 0’). SMC enrichment zones determined by ChIP-seq **(Figure S2**) are indicated by magenta bars.

Here, we investigate the collisions between the SMC complex and the replisome *in vivo*. We have taken advantage of the *B. subtilis* system, which much better tolerates the manipulations of SMC loading than other bacterial species ^25, 30, 31^. We have controlled SMC loading spatially by changing the location of SMC loading sites (*parS*), and temporally by inducing the expression of the SMC loader (ParB) at desired times. Combined with our ability to control the initiation and progression of DNA replication using mutants and drugs, we have assessed the fate of SMC complexes and the replisome when they collide in a head-on or a head-to-tail orientation. We tracked the changes in chromosome organization, SMC distribution, and DNA replication over time using chromosome conformation capture (Hi-C) assays, chromatin immunoprecipitation with deep sequencing (ChIP-seq) assays, and genomic profiling. Then, we combined experimental data with simulations of SMC translocation and replisome movement to determine whether SMCs disassociate, stall, continue translocating, or exhibit a combination of these activities when encountering the replisome.

## Results

### SMC distribution does not affect replication profile

We first investigated the effect of SMC occupancy on replisome progression. Since SMC is loaded by ParB binding to *parS*, strains containing different *parS* sites have varied SMC occupancy and chromosome folding patterns. We chose three strains: 1) the wild-type *B. subtilis* strain containing 9 *parS* sites, which had a gradient of SMC localization from origin to the terminus by ChIP-seq, and had a secondary diagonal and a star-shaped interaction pattern near the origin by Hi-C ^32^; 2) a strain lacking all 9 *parS* sites, which had uniform SMC distribution along the genome, and lacked a secondary diagonal on the Hi-C map ^25^; 3) a strain harboring a single *parS* site at -117°, which had SMC concentrated within one chromosome half, and whereby less than half of a chromosomal DNA was juxtaposed by SMC as seen by Hi-C ^25^ (**Figure 1B**). To understand whether the different SMC occupancy in these three strains affects replication progression, we performed whole-genome sequencing (WGS) coupled with marker frequency analysis (MFA). Our results showed that these three strains had near identical replication profiles, indicating that distributions and movement of SMC have no effect on replication progression in exponentially growing cells (**Figure 1B**).

### Synchronizing replisome progression

To directly test the effect of SMC translocation on the progression of the replication fork, we synchronized DNA replication using a *dnaB*(ts) allele ^32, 33^. Cells were first grown exponentially at a permissive temperature (30°C), then shifted to a restrictive temperature (42°C) for 45 min. This step allowed DNA replication to finish but inhibited new rounds of replication, arresting cells at G1. Next, cells were shifted down to 30°C to initiate replication synchronously. We incubated the cells at 30°C for only 15 min, then shifted them back to 42°C. This step allowed the current round of replication to progress, but prevented further initiation events, resulting in a single round of synchronized replication. We took samples at indicated time points and used MFA to analyze replication progression (**Figure 1C**). In all the experiments mentioned in this study, we used the same procedure for the replication synchronization.

The MFA plots showed that cells achieved reasonable synchrony for replication initiation, progression and termination (**Figure 1C**, blue lines). However, there were features indicating that the synchronization was not perfect (**Figure S1A**). First, at T = 0 before initiation, instead of a flat line, there was a shallow slope from the origin to the terminus (**Figure 1C** and **Figure S1A**, slope 1), indicating that a fraction of cells was not arrested at G1 but had pre-existing replisomes on the chromosome. Secondly, during replication, the peak height was lower than two folds of the baseline (**Figure S1A**, middle panel), indicating that not all the cells in the population had initiated replication, resulting in a small fraction of cells without replisomes. Thirdly, instead of a vertical line connecting the replicated region to the unreplicated region, there was a slope in the experimental profile (**Figure S1A**, slope 2), indicating that cells started replication at within a time window. Finally, after the synchronized round of replication was completed, the replication profile had a slope (**Figure S1A**, slope 3), indicating that replisomes were still present in a small fraction of cells.

To understand the replisome progression in our experiment, we performed simulations of replisome dynamics to reproduce the distribution of replisomes according to the MFA plots (see details in **Supplemental Methods**). We varied five parameters of replisome behavior including the fraction of cells containing replisomes before initiation (e.g. pre-existing replisomes), the average time for replisome to load to the origin, the speed of replisome movement at 30°C and 42°C, and the rate of spontaneous replisome stalling after replication. We identified one set of numbers that reproduced the MFA profiles for all the time points in this experiment (**Figures S1B** and **S1C),** as well as all the MFA plots in later figures (**Figure S1D-F**). Our simulations indicate that the fraction of pre-existing replisome was 18%, the speed of DNA replication at 42°C was 66 kb/min, and the calculated fraction of cells that did not initiate replication was ∼10%.

### SMC translocation does not affect replisome progression

To directly test whether SMC translocation affects replisome progression, we induced SMC translocation in replication-synchronized cells and monitored the difference in fork progression with and without translocating SMC (**Figure 1C**, red and blue lines). The location of SMC loading was specified by a single *parS* site at -59°, and timing of SMC loading was controlled by an IPTG-inducible *parB* as the sole source of ParB protein. We added IPTG at the onset of replication initiation. ChIP-seq results showed that SMC was loaded and enriched progressively as expected (**Figure S2**). Judged by the SMC enrichment zone (**Figure 1C**, magenta bars determined by **Figure S2**) and replisome progression (**Figure 1C**), SMC-replisome collisions occurred from 15 min to 50 min. Nonetheless, in the presence of SMC, the replication progression profiles overlapped with ones without SMC (**Figure 1C**, red and blue curves). These results indicate that SMC translocation does not affect replication progression.

### SMC translocation is affected by replisomes

Conversely, we investigated the impacts of replisome movement on SMC translocation (**Figure 2**). We used a strain that retained the endogenous *parS* at -27° but lacked the other eight *parS* sites. This strain also contained a *dnaB*(ts) allele to allow for synchronization of DNA replication using the procedure described above (**Figure 2A**, MFA plots). At T = 0, cells were arrested at G1 as indicated by the MFA plot. The Hi-C map had a secondary diagonal emanating from -27°, showing that the DNA regions flanking the *parS* were juxtaposed (**Figure 2A** and **2B**, T = 0, line 1). From 0 min to 15 min, replication initiated and progressed toward the *parS* site. The replisome collided head-on with SMCs. We observed three new features on the Hi-C maps. First, starting at -27°, the secondary diagonal curved downward (**Figure 2A**, arc 1), indicating that SMCs colliding with the replisome moved slower than the ones translocating toward the terminus. Secondly, there was a gap on the secondary diagonal (**Figure 2A**, gap), which could be generated by SMC slowing down or dissociation when colliding with the replisome (**Figure 2B**, T = 10’-15’). Thirdly, the remaining portion of the secondary diagonal towards the terminus region (**Figure 2A**, T = 10’-15’, line 1) maintained the curvature seen at T = 0, indicating that SMCs running ahead of replication forks were unaffected by the replication. It is important to note that, arc 1 extended beyond the replisome, which became more evident at later time points. This observation indicates that some SMC bypassed the replisome and continued translocation.

**Figure 2.**
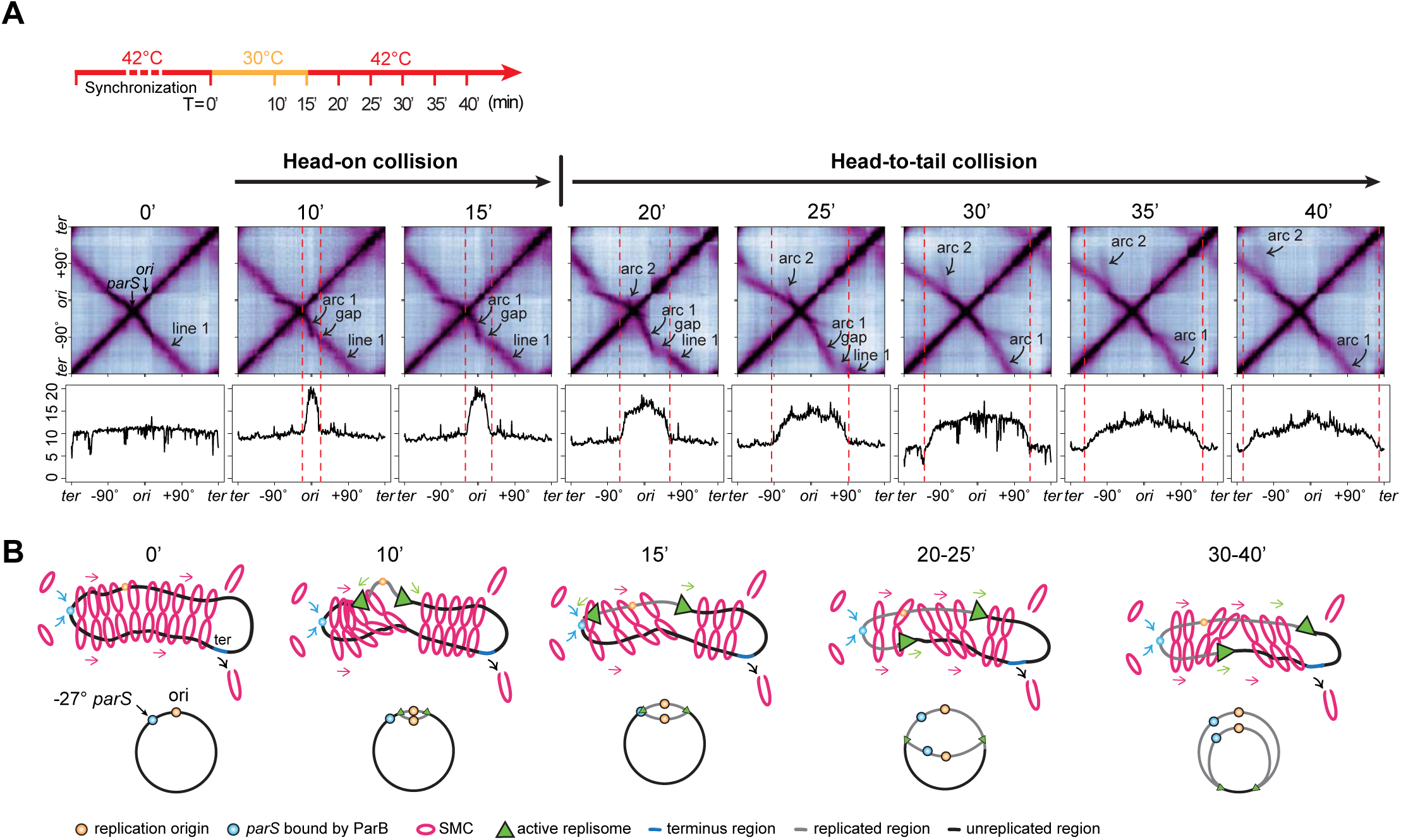
SMC translocation is affected by replisomes. **(A)** Time-course Hi-C maps and MFA plots of a strain containing a single *parS* site at - 27° (BWX5230). G1-arrested cells were grown as indicated in the timeline (top row). T = 0’ indicates the onset of replication initiation. Samples were collected at indicated time points. **(B)** Schematics depicting major SMC-replisome engagements at the indicated time points (top row). For simplicity, only one copy of the replicated region is shown (gray). Replisome progression is indicated on the bottom row.

At 20-35 min, the replication fork moved beyond the *parS* site as indicated by MFA. The leftward replisome collided with SMCs in a head-to-tail manner (**Figure 2B**, T=20’-25’). The three features discussed above remained despite some minor changes: 1) arc 1 continued to grow, and the growing tip of arc 1 still curved down near the replication fork; 2) the gap moved; 3) line 1 retreated towards the terminus. Additionally, we found that the central portion of arc 1, which was close to the *parS*, flattened out to similar curvature as seen at T = 0, indicating that newly loaded SMCs were not affected by the replisome far ahead. In addition to these three features, we found a new smaller arc (arc 2) emanating from the *parS* (−27°). The arc curved up and tracked immediately behind the replication fork. The shape of arc 2 indicates that the SMC subunit heading towards the terminus region traveled more slowly than the subunit heading towards the replication origin. This is consistent with the idea that SMCs loaded behind the replisome reached the replisome; one SMC subunit collided with the replisome in a head-to-tail manner and was slowed down by the replisome, whereas the other SMC subunit translocating towards the origin region was not affected.

These results demonstrate that when encountering with the replisome, SMC movement was altered. The Hi-C maps show hints of SMC paused by the replisome **(Figure 2A**, arc 1 and gap), and SMC bypassing the replisome (**Figure 2A**, 10-30 min, arc 1). It is also possible that some SMCs dissociated after the collisions. Given the continuous loading of SMCs, the continuous movement of replisomes, and concurrence of head-on and head-to-tail collisions between SMC and the replisome, it is difficult to quantify the consequences of SMC-replisome collisions in this experimental setup. In our next sections, we set up simpler scenarios of SMC-replisome collisions to dissect the rules of engagement between SMC and the replisome.

### Engineering an *in vivo* system for SMC-replisome collisions

To better resolve the effect of the replisome on SMC translocation, we designed a system to exert both temporal and spatial controls over the collisions between SMC and the replisome on the chromosome. For SMC, we inserted a single *parS* site at different genome positions to control SMC loading spatially and an IPTG-inducible *parB* to control the timing of loading. For the replisome, we used the *dnaB*(ts) allele to control the timing of replication initiation. We asked whether we could pause replisomes at specific locations to better manage the collision site. HPUra is a specific inhibitor of DNA polymerase PolC in *B. subtilis*, which has been shown to stall replication but retain the replisome at the replication fork ^34, 35, 36^. To test the effectiveness of HPUra in our experimental system, we combined HPUra treatment with temperature shift experiments using a *dnaB*(ts) strain. We monitored replication progression using MFA and examined replisome integrity using fluorescence microscopy of DnaX-YFP ^32^ (**Figure 3**).

**Figure 3.**
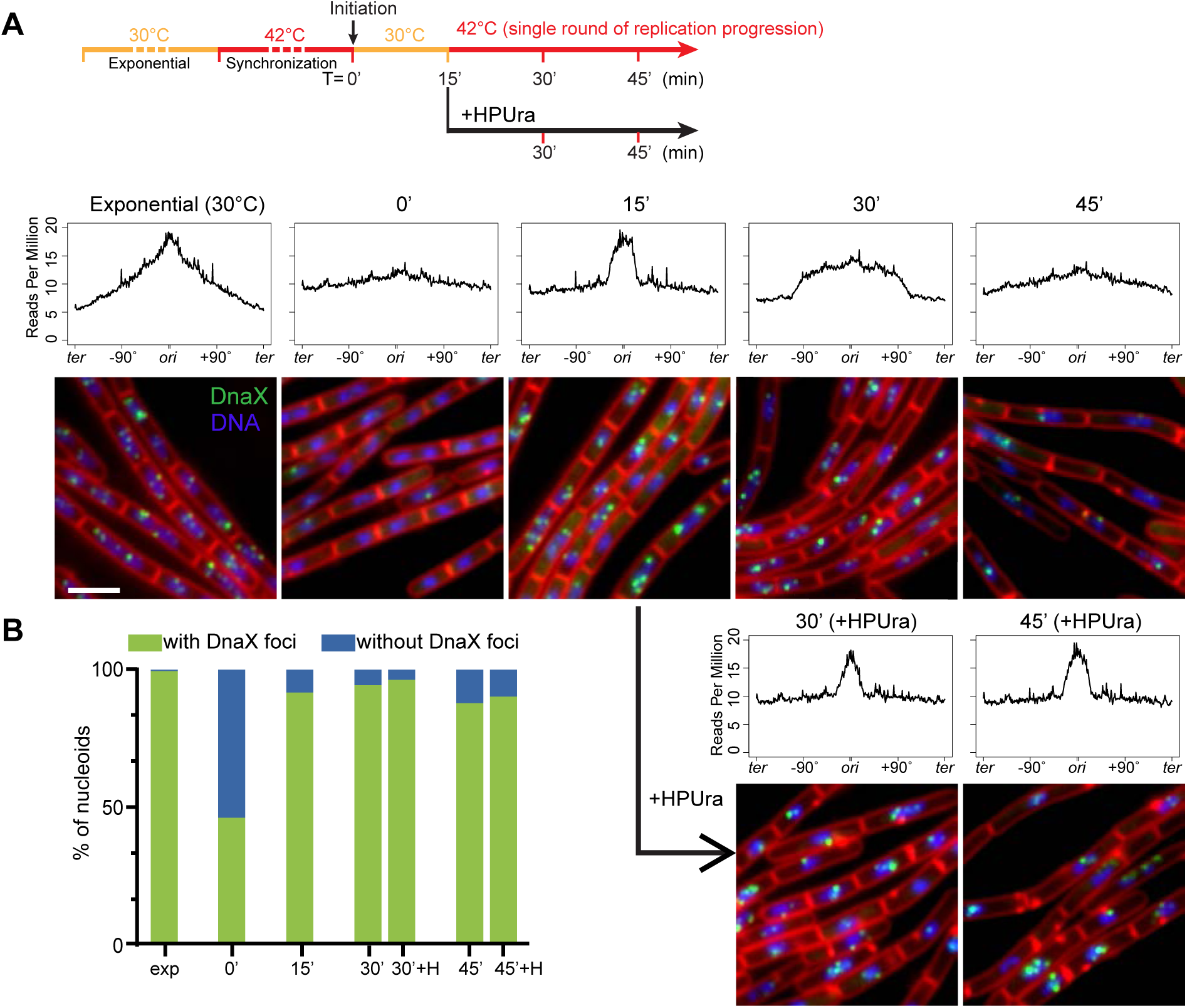
HPUra stalls replication but does not affect replisome assembly. See also Figure S3. **(A)** As shown in the experiment timeline (top row), a *dnaB*(ts) strain (BWX4310) was grown exponentially at 30°C for 2 h, shifted to 42°C for 45 min to synchronize DNA replication. At T = 0’, cells were shifted to 30°C for 15 min to initiate replication then back to 42°C to allow a single round of replication. HPUra was added at T = 15’. Middle panel: MFA plots are shown for indicated time points. Bottom panel: representative images of a *dnaB*(ts) strain (BWX2533) grown as described above. The images contain DAPI-stained nucleoid (blue), FM-4-64-stained membrane (red) and YFP-tagged DnaX (green). Scale bar represents 2 μm. **(B)** Analysis of imaging results in (**A**). Nucleoids with or without DnaX foci were quantified at indicated time points and the percentages were plotted.

Cells were synchronized as described above (**Figure 3A**, timeline). At 15 min after replication initiation, the cell cultures were treated with or without HPUra. In absence of HPUra, replication progressed as expected (**Figure 3A**, MFA plots). In the culture treated with HPUra, the DNA content remained unchanged over time (**Figure 3A**). These results indicate that HPUra stalled replication nearly immediately, and the DNA remained intact without degradation.

To test whether the replisome remains bound to the replication forks after HPUra treatment, we visualized the DnaX-YFP foci using fluorescence microscopy (**Figure 3A**, micrographs). In the exponential growth phase, DnaX-YFP foci were detected in 99.4% of nucleoids, indicating active replication. After G1 arrest, we observed that 46.2% of nucleoids contained the replisome foci, much higher than the 18% of cells with existing replisomes as indicated by MFA (**Figure S1B**). The higher percentage was derived from a portion of *dnaB*(ts) cells that had initiated replication during the sample preparation for the imaging which was performed at room temperature. At 15 min after initiation, we found the replisome foci in 91.7% of nucleoids, consistent with our earlier estimation that ∼10% of cells did not initiate replication measured by MFA results (**Figure 1C** and **Figure S1**). Over time, the percentage of nucleoids containing the foci dropped slightly due to the cells completing replication. In the presence of HPUra, the percentage of cells containing the replisome foci remained largely unchanged over time (**Figures 3A** and **3B**). These results indicate that the addition of HPUra stalled the replication effectively and the replisomes remained at the replication fork, consistent with previous reports ^35, 36^.

To test whether HPUra has an off-target effect on SMC translocation, we performed Hi-C on *dnaB*(ts) cells that were constantly arrested at G1 (**Figure S3A**). At 15 min after ParB induction, ∼730 kb of DNA from each of the left and right arms were juxtaposed by SMC. After another 15 min, with or without HPUra, DNA juxtaposition proceeded to the same distance (**Figure S3A**). Therefore, in the absence of active DNA replication, HPUra does not affect the SMC translocation; the effect of HPUra on SMC movement reported in the following sections are due to HPUra’s effect on the replisome.

### Head-to-tail collisions between SMC and the replisome (two sided)

In *B. subtilis*, SMC translocation is faster than DNA replication: in exponentially growing cells, SMC translocation is at ∼50 kb per minute while the replisome progresses at ∼40 kb per minute ^25, 37^. In our current experimental setup at 42°C, SMC translocates at ∼71 kb per minute (**Figure S3B**) while the replisome progresses at ∼66 kb per minute (**Figure S1B**). Thus, SMC that loads behind the replisome may catch up and collide with the replisome in a head-to-tail orientation. Below, we set up two experiments to examine this type of collision.

Many bacteria have *parS* sites that are very close to replication origin ^29^. *B. subtilis* has five origin-proximal *parS* sites at -1°, -3°, -4°, -5° and +4°, respectively (**Figure 1A**). After replisomes travel past these sites, SMCs loaded at these sites will track behind both replisomes, catch up with them, and collide into both replisomes at about the same time. To understand the consequences of such two-sided head-to-tail collisions, we used a strain containing a single *parS* at the endogenous -1° site. In a control condition in which the replisome was far ahead of the SMC, there was little chance of SMC-replisome collision, and the whole chromosome arms gradually juxtaposed at ∼71 kb per minute (**Figure 4A**). To set up an experiment for SMC-replisome collision, we allowed synchronous replication initiation for 15 min, then added HPUra to pause replication forks (**Figure 4B**, MFA plots). At the same time, we induced expression of ParB to load SMC, then monitored DNA juxtaposition over time (**Figure 4B**, Hi-C plots). Comparing with the control, the secondary diagonal in the collision experiment had three major features: 1) The Hi-C intensity had a sharp drop beyond the replisome, suggesting that the majority of SMCs did not bypass the replisome, but were either blocked or removed by the replisome; 2) Although the intensity was faint, the secondary diagonal extended beyond the replisome, suggesting that some SMCs were able to bypass the replisome and continue zipping up the arms; 3) The endpoints of DNA juxtaposition lagged behind those in the control by about 200 kb on each arm. Using the measured extrusion rate of ∼71 kb per minute, we inferred that a small fraction of SMCs paused for about three minutes before bypassing the replisome.

**Figure 4.**
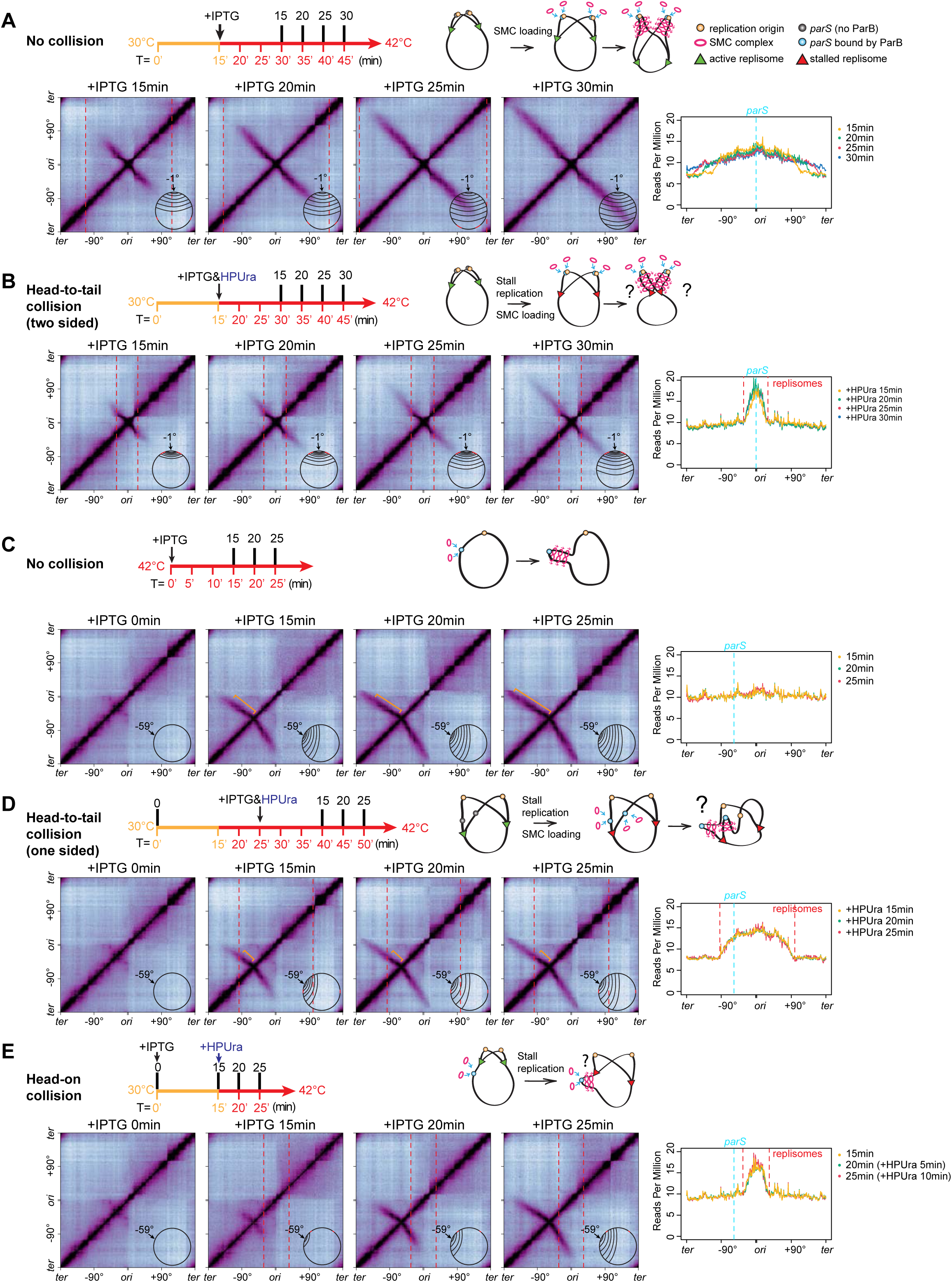
Head-to-tail and head-on collisions between SMC and stalled replisomes. See also Figure S3. Five experiments to set up different SMC-replisome collisions by controlling SMC loading (using IPTG-inducible ParB) and replication stalling (using HPUra) in cells synchronized for DNA replication (using a *dnaB*(ts) allele). The timeline for each experiment is shown on the top panels. Numbers below the timeline indicate synchronization status, in which T = 0’ is the onset of replication initiation; numbers above the timeline indicate the time after IPTG/ HPura addition) as indicated. Schematics of experimental setups are shown on the top right panel. **(A)** A control experiment with no SMC-replisome collisions, in which replisome is far ahead of SMC. A *dnaB*(ts) strain containing IPTG-inducible ParB and a single *parS* at - 1°(BWX4310) was grown as indicated by the timeline. Hi-C contact maps were generated from cells at 15 min, 20 min, 25 min and 30 min after IPTG addition. Schematics of juxtaposed region are superimposed on the Hi-C maps. The positions of replication forks determined by MFA plots are indicated by red dotted lines. MFA plots for all indicated time points are plotted in a single graph (bottom right), on which the *parS* site is indicated by a cyan dashed line. **(B)** Two-sided head-to-tail collision in which both SMC motors encounter stalled replisomes. In the same strain shown in (**A**), HPUra was added to stall replication at 15 min after replication initiation. At the same time, IPTG was added to induce SMC loading. **(C)** A control experiment with no SMC-replisome collision using a *dnaB*(ts) strain containing IPTG-inducible ParB and a single *parS* at -59° (BWX5297). This control was set up by growing G1-arrested cells at 42°C constantly (see timeline in the top row). Hi-C contact maps and the MFA plots are shown for samples at 0 min, 15 min, 20 min, 25 min after IPTG addition. The regions with Hi-C interaction score more than 0.008 are indicated by orange bracket-like marks. **(D)** One-sided head-to-tail collision in which one motor of SMC complex encounters a stalled replisome. In the same strain shown in (**C**), after the leftward replication fork moved beyond the *parS* site, the replisomes were stalled by the addition of HPUra. At the same time, SMC loading was induced by IPTG. The regions with Hi-C interaction score more than 0.008 indicated by orange bracket-like marks. **(E)** Head-on collision between SMC and the replisome. In the same strain shown in (**C**), the replisomes were stalled before colliding with SMC. Hi-C contact maps and the MFA plots are shown from samples at 0 min, 15 min, 20min, and 25 min after IPTG induction.

### Head-to-tail collisions between SMC and the replisome (one sided)

Many bacterial genomes contain *parS* sites far away from the origin ^29^. *B. subtilis* has four *ori*-distal *parS* sites at +17°, +42°, +91°, -27°, respectively (**Figure 1A**). After a replisome passes these sites, SMCs loaded behind the replisome may catch up with only one replisome, generating a one-sided head-to-tail collision. To investigate this situation, we engineered a single *parS* site at -59° site (**Figures 4C** and **4D**). In the control experiment, SMC loading was induced in cells arrested in G1 without replication (**Figure 4C**, MFA plots). The time-course Hi-C experiment shows that DNA juxtaposition proceeded at a constant speed, ∼71 kb per minute towards the terminus and ∼49 kb per minute towards the origin, and the zipping was completed after 25 min of the induction (**Figure 4C**, Hi-C plots).

To set up one-sided head-to-tail collision, we initiated replication and allowed the replisome to progress beyond the *parS* site (**Figure 4D**, MFA plots). We added HPUra to pause replication and added IPTG to induce the SMC loading (**Figure 4D**). In the Hi-C time course, compared with the control experiment without replisome present (**Figure 4C**), the secondary diagonal in the collision experiment had the same major features observed in **Figure 4B**: DNA juxtaposition extended beyond the collision site, but the intensity was much lower than that in the control, suggesting that the majority of SMCs were removed or paused by the replisome, and some SMCs bypassed the replisome. Moreover, the distance of DNA juxtaposition towards terminus was ∼ 133 kb less in **Figure 4D** than in **4C.** Using the terminus-proximal extrusion rate of ∼71 kb/min, we inferred that bypassing happened after a about two-minute delay in this one-sided head-to-tail collision. Consistent with the Hi-C results, ChIP-seq analyses reveal that SMCs enrichments decreased beyond the collision site compared with the control (**Figure S3C**).

### Head-on collisions between SMC and the replisome

Since all the *parS* sites have some distance from *oriC*, as the replisome travels from origin towards the terminus, SMC loaded at the *parS* will collide with the replisome head-on. To explore the consequence of head-on SMC-replisome collisions in a simple setup, we used the strain containing a single *parS* site at -59°. We induced SMC loading at the onset of replication initiation and added HPUra at 15 min after replication initiation, which was before the replisome encountered SMC (**Figure 4E**). Despite some delay, the DNA juxtaposition extended beyond the replisome, indicating that some SMCs bypassed the replisome. However, the intensity beyond the collision point was lower than that of the control (**Figure 4C**), indicating that some SMCs were unloaded or paused at the collision site.

### Framework of simulations

Next, we used simulations to understand what rules of engagement between SMC and the replisome generated the observed Hi-C and ChIP-seq results. We first simulated replisome dynamics to reproduce the experimental replication profiles (**Figure S1**, also see details in **Supplemental Methods**). For SMC dynamics, we adapted a framework based on previous measurements and simulations ^38, 39, 40^ and included parameters specific to this study (see details in **Supplemental Methods**). Briefly, we set the number of SMC complexes (loop-extruding unit) to be 40 per chromosome. We considered that one loop-extruding unit contained two independent extrusion motors translocating away from the loading site. If translocation by one motor was blocked by an encounter with the replisome, the other continued its extrusion. SMC complexes were allowed to load anywhere on the chromosome, with a loading preference at the *parS* site, so that most loaded at *parS*. In addition, we determined two parameters that were specific for this study’s experimental conditions: we found SMC’s spontaneous disassociation rate at 42°C (**Figure S4**, also see details in **Supplemental Methods**), and measured the SMC translocation rates at 42°C as 71.5 ± 2 kb/min per motor subunit (i.e. for a total rate of loop growth of 143±4 kb/min) (**Figure S3B**).

When SMC reached a replisome, we first considered three outcomes ^20^: blocking in which the collided SMC subunit was stopped by the replisome; unloading in which the whole SMC complex was removed by the replisome; and bypassing in which SMC traversed the replisome and continued translocating. Given the indications from experiments, we next extended these simple models by allowing SMC to pause in a blocked state for a varying amount of time before unloading or bypassing. We simulated SMC dynamics and calculated SMC occupancy on the chromosome to compare with the experimental ChIP-seq results. Using the SMC chromosomal positions, we generated simulated contact frequency maps to compare with experimental Hi-C results (see details in **Supplemental Methods**).

### Determining the rules for head-to-tail SMC-replisome collisions

We studied the rules of engagement for SMC and the replisome in the head-to-tail collision shown in **Figure 4D**. We used the experimental data at 25 min following IPTG treatment (**Figure 5A**) and first explored the three basic rules of engagement: bypassing only, blocking only, and unloading only (**Figures 5B-D**). In each model, the simulated SMC ChIP enrichment profiles (**Figures 5B-D**, orange lines) and Hi-C maps (**Figures 5B-D**, bottom panels) showed specific features that were different from the experimental data (**Figure 5A**).

**Figure 5.**
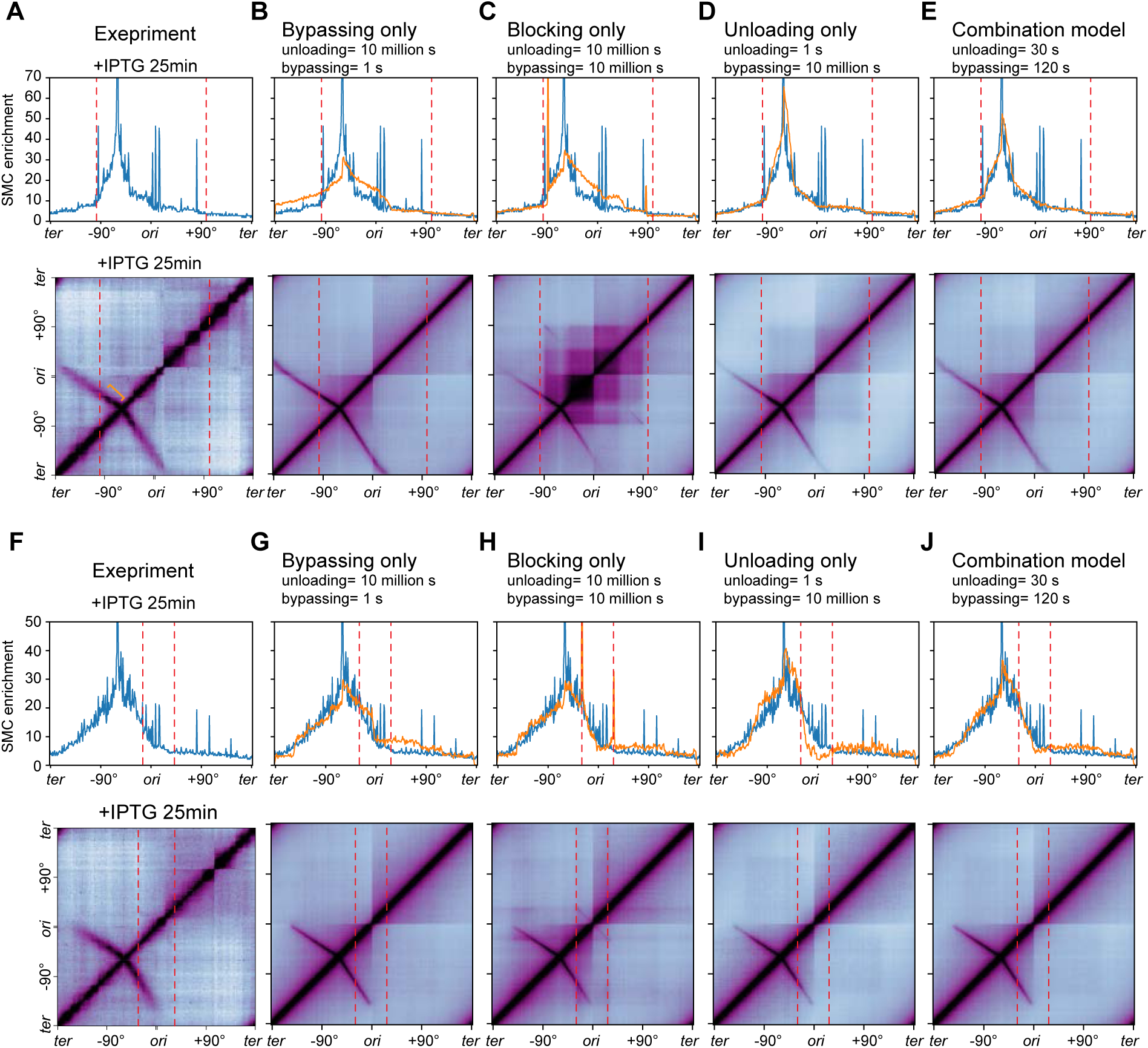
Simulation results for head-to-tail and head-on SMC-replisome collisions. See also Figures S5 and S6. **(A)** Experimental data for head-to-tail collision shown in **Figure 4D** (+IPTG 25 min). Top: anti-SMC ChIP-seq was filtered as described in Methods and plotted as reads per million in 10-kb bins. Bottom: Hi-C contact map. The positions of replication forks are indicated with red dashed lines. **(B-E)** Top: simulated SMC distributions (orange curves) plotted on top of experimental ChIP-seq result (blue curves). Bottom: simulated Hi-C maps. Four models are shown respectively for bypassing-only (**B**), blocking-only (**C**), unloading-only (**D**), and the combination model (**E**). The complete parameter sweep is shown in **Figure S5**. **(F)** Experimental data for head-on collision shown in **Figure 4E** (+IPTG 25 min). Top: anti-SMC ChIP-seq was filtered as described in Methods and plotted as reads per million in 10-kb bins. Bottom: Hi-C contact map. **(G-J)** Top: simulated SMC distributions (orange curves) plotted on top of experimental ChIP-seq result (blue curves). Bottom: simulated Hi-C maps. Four models are shown respectively for bypassing-only (**G**), blocking-only (**H**), unloading-only (**I**), and the combination model (**J**). The complete parameter sweep is shown in **Figure S6**.

In the bypassing-only model, the simulated SMC enrichment beyond the collision site was much higher than the experimental ChIP result (**Figure 5B**, top panel). The simulated Hi-C map had the DNA regions flanking the *parS* site fully juxtaposed, with intensities very similar to the control which had no collision between SMC and the replisome (the simulated Hi-C map in **Figure 5B** compared with the experimental Hi-C map at +IPTG 25min in **Figure 4C**). These results indicate that the bypassing-only model allows too many SMCs to go beyond the replisome.

In the blocking-only model, simulated SMC occupancy showed a sharp peak at the collision site (near -90°), which was absent from experimental ChIP-seq data. Additionally, SMC enrichment on the right arm was much higher than the experimental data (**Figure 5C**, top panel). The simulated Hi-C map showed an intense “square” of interactions at the center of the map (**Figure 5C**, bottom panel), which was absent in the experimental Hi-C map (**Figure 5A**, bottom panel). Therefore, we ruled out the blocking-only model.

The unloading-only model exhibited a better match to the experimental data, but the ChIP signal near the loading site was higher than the experimental data (**Figure 5D**, top panel). The simulated Hi-C interaction frequency beyond the collision site diminished much faster than the experimental Hi-C (**Figure 5D**, bottom panel).

Our results indicate that the single rule of engagement (bypassing, blocking, or unloading) did not fully reproduce the experimental data. We next explored a combination of these engagement modes to achieve a better fit (**Figure S5**). We assumed that when SMC collided into a replisome, the collided SMC motor was blocked, then it either bypassed the replisome or unloaded from the chromosome at specific rates (see details in **Supplemental Methods**). We generated simulated SMC occupancy profiles across a broad range of bypassing and unloading times to compare with experimental data (**Figure S5A)**. Intuitively, when the bypassing time was short and the unloading time was long, the simulated result resembled the bypassing-only model (**Figures 5B** and **S5A**); when the both the bypassing and unloading times were long, the result resembled the blocking-only model (**Figures 5C** and **S5A**); when the unloading time was short and the bypassing time was long, the result resembled the unloading-only model (**Figure 5D** and **S5A**). To find the optimal model matching the experimental data, we performed goodness-of-fit analysis (**Figure S4B,** also see details in **Supplemental Methods**) and visual inspection of the overall shape of SMC enrichment profiles to assess the similarity between the simulations and the experiment. We found that if the unloading was more than two-fold faster than the bypassing, the simulation produced a better fit than otherwise (**Figure S5B**, 1:2 ratio line). A few combinations produced a very good fit (**Figure S5B,** black asterisks). Since the experimental Hi-C analysis estimated the bypassing time to be about two minutes for this one-sided head-to-tail collision (**Figures 4C** and **4D**; **Figure S5B,** white rectangle), the best combination in the range was unloading at 30 s and bypassing at 120 s (**Figure S5B**, black check mark), which produced simulated ChIP and Hi-C results better matching the experimental data than models involving the single rule of engagement (**Figures 5E** compared with **5B-D**).

### Determining the rules of SMC-replisome head-on collisions

Next, we investigated the rules of engagement for SMC and the replisome in the head-on collision shown in **Figure 4E**. We simulated SMC occupancy profiles and Hi-C results at 25 min after IPTG induction (**Figures 5F-J**). In a trend similar to the head-to-tail simulations, simple models of bypassing, blocking, or unloading did not reproduce the experimental results (**Figures 5F-I**). Next, we swept a wide range of unloading and bypassing times (**Figure S6A**). Goodness-of-fit analysis and visual inspection identified four combinations that produced much better fits than each of the simple models (**Figure S6B,** black asterisks). Among these combinations was the same rule of engagement seen for head-to-tail collision (unloading=30 s and bypassing=120 s) (**Figures 5J** and **S6B**, black check mark).

Although our simulations did not narrow the unloading and bypassing times to one specific set of numbers, we found that unloading is the major rule and bypassing is less frequent. Although it is possible that SMC-replisome head-on collisions and head-to-tail collisions use different unloading and bypassing rates, we found certain combinations, such as unloading at 30 s and bypassing at 120 s, can explain both types of collisions. This set of numbers means that, when SMC encounters the replisome, on average, it pauses for 24 seconds, then with 80% chance unloads from the chromosome, and with 20% chance, it bypasses the replisome and continues translocation.

### SMC colliding with a moving replisome

In experiments shown in **Figure 4** and simulations in **Figure 5**, we used stalled replisomes, which simplified the collision scenarios but lacked replisome movement and activity. Our earlier results showed that the replisome proceeded normally regardless of collisions with SMCs (**Figure 1C**). Thus, we hypothesized that during the time when SMC was blocked by a moving replisome, the replisome would push SMC at head-on collisions and would retard SMC to the same speed as the replisome at head-to-tail collisions. To test this hypothesis and understand whether the rules of engagement determined using stalled replisomes apply to unperturbed replisomes, we performed two SMC-replisome collision experiments with moving replisomes.

In the first experiment, we used the same strain as in **Figures 4D** and **4E**. We induced SMC loading upon replication initiation and allowed the replisomes to progress through the cell cycle (**Figure 6A**). MFA and Hi-C results showed that at T = 15-20 min, SMCs collided with the moving replisome head-on. Compared with experiments containing the stalled replisome, SMC colliding with a moving replisome had stronger features: on the Hi-C maps, DNA juxtaposition progressed slower, and the secondary diagonal had a stronger downward tilt (compare **Figure 4E** and **Figure 6A** at 20 min**; Figures S7A** and **B**); on ChIP-seq plots, SMC enrichment had a slower progression and stronger buildup over the collision zone (**Figures S7C** and **D**, black arrows), although SMC enrichment on terminus-proximal side was unaffected. These results are consistent with the notion that the collided SMC motor was blocked and pushed back by the moving replisome, while the unblocked SMC motor continued translocating. Despite this block and pushback, DNA juxtaposition eventually extended beyond the replisome albeit at lower intensity, indicating that some SMC bypassed the moving replisome. After the events of head-on collisions, the intensity of the secondary diagonal was reduced (compare 25 min with 20 min in **Figure 6A**; compare moving replisome with stalled replisome in **Figure S7A** and **B**, 25 min), suggesting that some SMCs were unloaded from the chromosome upon head-on collisions. After the replisome passed the -59° *parS* site (**Figure 6A**, T = 25-40 min), SMCs loaded at *parS* caught up and collided with the replisome in a head-to-tail orientation. The emergence of a new arc on the Hi-C map (**Figure 6A**, 30 min black arrow) suggests that some SMCs were slowed down when colliding head-to-tail with the moving replisome.

**Figure 6.**
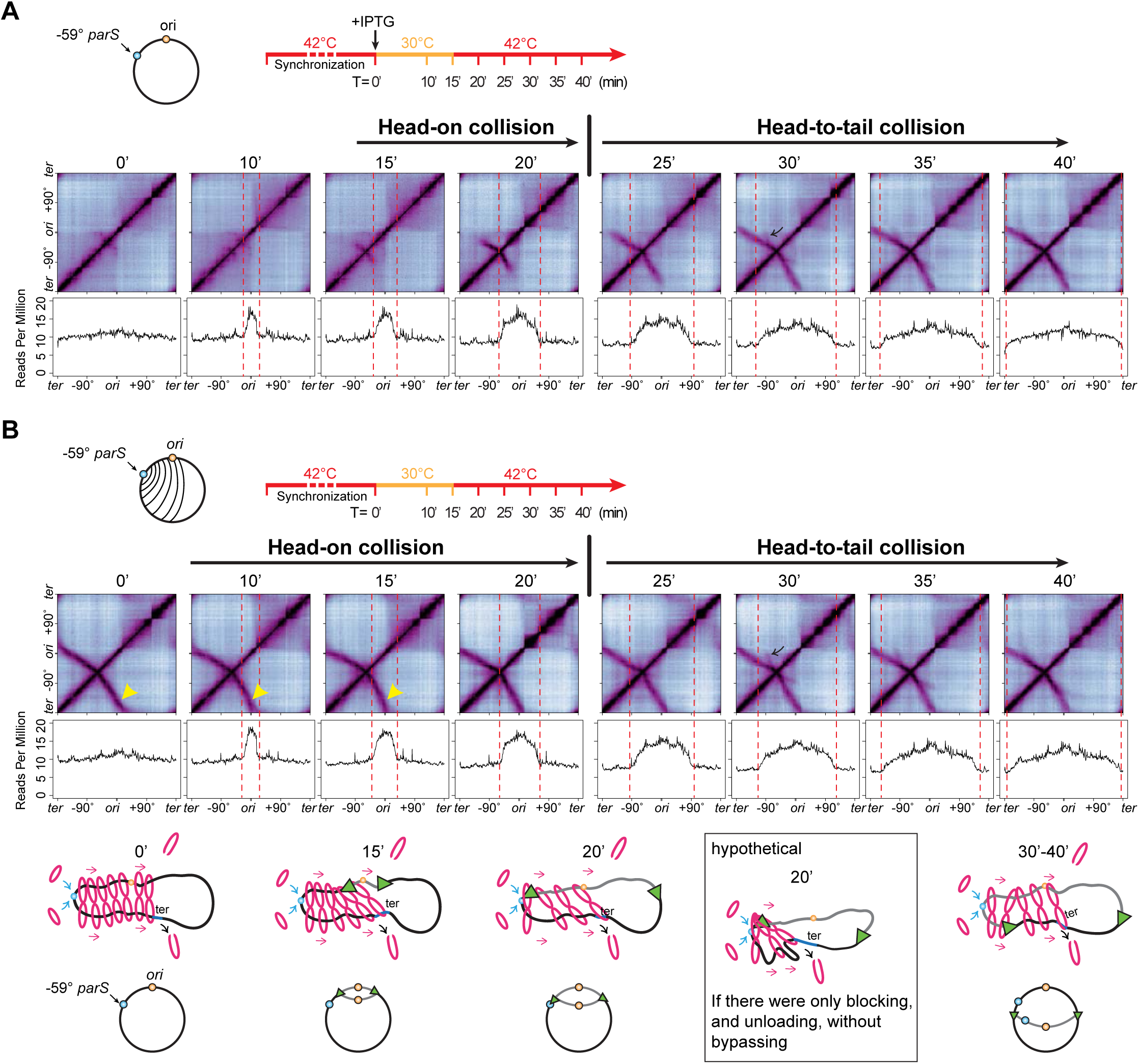
The collisions between SMCs and moving replisomes. See also Figure S7. **(A)** Hi-C maps and MFA plots generated from a time course experiment using a *dnaB*(ts) strain containing IPTG-inducible ParB and a single *parS* at -59° (BWX5297), which is the same strain used in Figure 4C-E. SMC loading was induced at the onset of replication initiation. In this experiment, replisomes were allowed to progress through the cell cycle, in contrast to **Figure 4D-E** which stalled replisomes using HPura, **(B)** Hi-C maps and MFA plots generated from a time course experiment using of a *dnaB*(ts) strain containing a single *parS* at -59° *parS* (BWX5529). In this strain, SMC is loaded constantly, in contrast to IPTG-induced SMC loading seen in (**A**). Schematics depicting SMC translocation at indicated time points.

In a second experiment, we used a strain with continuous SMC loading at the -59° *parS* site (**Figure 6B**). This allowed SMC to pre-load on the chromosome, increasing the frequency of SMC-replisome collisions, particularly head-on encounters (T = 0-20 min). Hi-C analyses revealed evidence of blocking, bypassing, and unloading. First, the secondary diagonal showed a pronounced downward curve at 10-15 min (**Figure 6B**, compare with 0 min, yellow carets), indicating that the moving replisome blocked and pushed SMC (**Figure 6B**, schematics 0 min and 15 min). Second, from 10-20 min, despite the replisome traversing the region between the origin and *parS*, this left arm region remained juxtaposed with the terminus-proximal region (**Figure 6B**, schematic 20 min). This persistent interaction indicates that some SMCs bypassed the replisome, as blocking and unloading alone would preclude such juxtaposition (**Figure 6B**, schematic inset). Finally, the fainter secondary diagonal generated by these bypassing SMCs (**Figure 6B**, compare 10-25 min with 0 min) suggests that a fraction of SMCs was unloaded upon collision.

Our experiments collectively demonstrate that SMC complexes, upon colliding with moving replisomes, undergo momentary blocking followed by either unloading or bypassing. This set of rules of engagement can also explain the results in our earlier experiment in **Figure 2**, in which we used a strain containing a -27° *parS* with continuous loading of SMC and moving replisomes. Finally, it is noteworthy that in three different time-course experiments with the moving replisomes (**Figures 2, 6A**, and **6B**), the replication profiles were similar regardless of the *parS* location and the timing of SMC loading, reinforcing the notion that SMC translocation does not affect replisome progression.

## Discussion

SMC complexes are major chromosome organizers in all domains of life. They load on the DNA and extrude DNA loops up to millions of base pairs in size, during which they encounter various DNA-bound molecules on the crowded chromatin fiber. Except for specific regulators, such as CTCF which blocks cohesion movement ^41, 42, 43, 44^ and the bacterial XerD protein which unloads SMC at the terminus region ^45^, SMC can bypass many molecules including nucleosomes ^46^, RNA polymerases ^39, 47^, nucleoid-associated proteins ^48^ and other SMC molecules ^38, 49, 50^. The ability for SMC to bypass barriers not only promote chromosome compaction and segregation, but also facilitate the trafficking of other factors along the chromosome ^40, 51^ ^52^. One important molecular machine that SMCs frequently encounter is the replisome. The *in vivo* consequence of their collisions and how these collisions affect DNA replication and chromosome compaction are unknown. Here, taking advantage of well-controlled SMC loading and replisome progression in *B. subtilis*, we determine the *in vivo* consequences of collisions between SMC and the replisome, and investigate the effects of SMC-replisome collisions on DNA replication and chromosome folding. Our experiments and simulations demonstrate that the progression of DNA replication is unaffected by SMC-mediated loop extrusion. By contrast, replisomes restrict loop-extrusion by blocking then unloading SMC complexes (with ∼80% chance). However, occasionally (with ∼20% chance) SMC can bypass the replisome and continue translocating.

### Head-on collision vs head-to-tail collision

Our experiments with moving replisomes (**Figures 2 and 7**) indicate that for both head-on and head-to-tail SMC-replisome collisions, a combination of blocking, unloading and bypassing must happen to generate the observed results. Our simulations show that unloading is more frequent than bypassing, and a combination of these modes of engagement (unloading=30 s and bypassing=120 s) (**Figure S5B and S6B**) can reproduce the prominent features of the experimental results for both head-on and head-to-tail collisions. Nevertheless, since the front and back of the replication fork have very different properties, for instance, at the front of the replisome there are positive supercoils, and behind the replisome there are single stranded DNAs and their binding proteins, Okazaki fragments, precatenated sister chromatids, and more, it is possible that the two types of collisions have slightly different rates for unloading and bypassing, which were not resolvable in our experiments.

**Figure 7.**
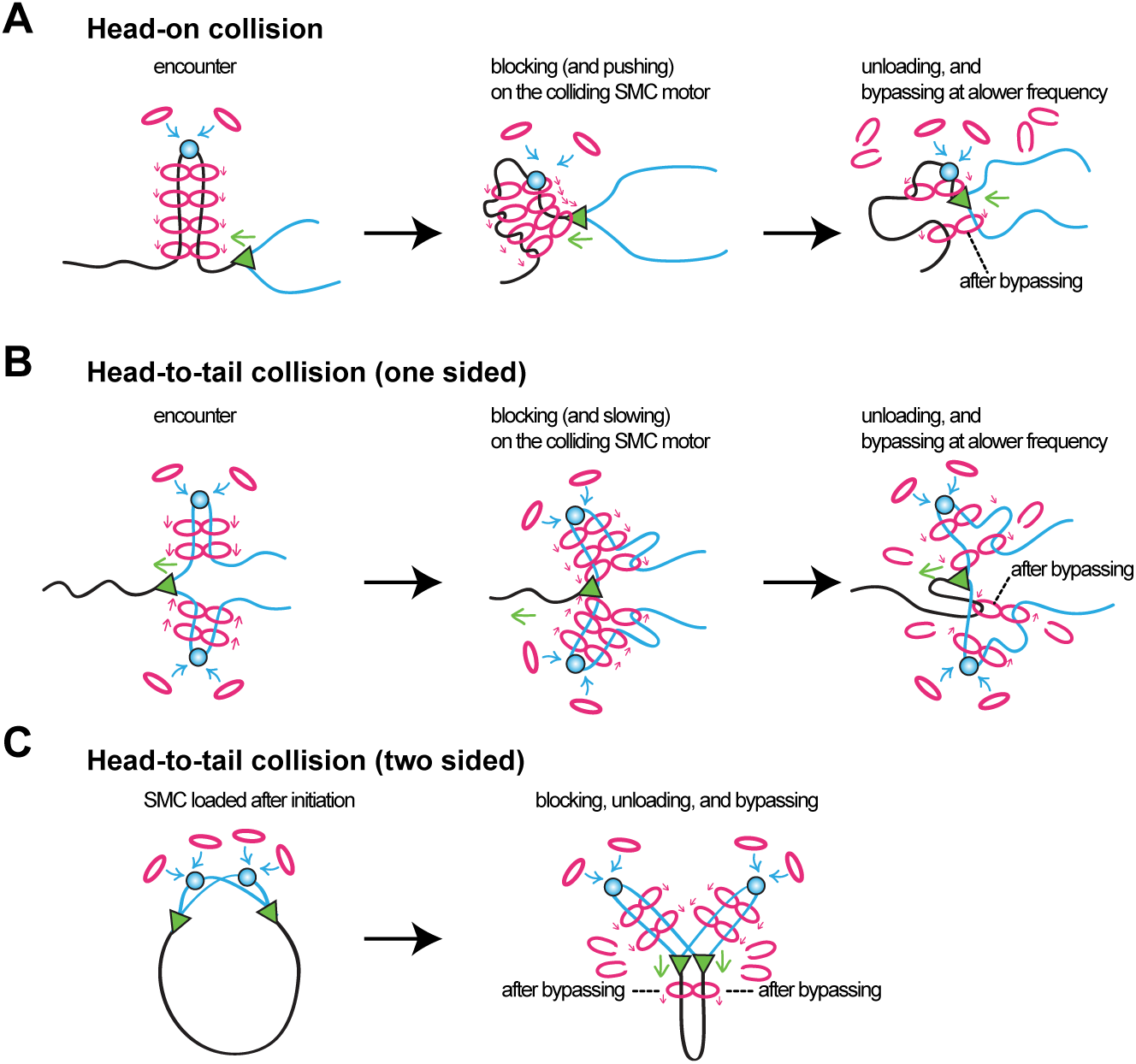
Schematic models for the consequence of SMC-replisome collisions. Models for SMC-replisome interaction for head-on collision (**A**), head-to-tail collision (**B**), and two-sided head-to-tail collision (**C**).

### SMC bypassing the replisome

Although the probability of SMC to bypass the replisome is low (∼20% or less), it is clear that bypassing is happening, as evidenced by DNA juxtaposition extending beyond the replisome (**Figures 2, 4**, and **7**). SMC bypassing the replisome could hinder chromosome segregation because this allows SMC to tether newly replicated DNA with unreplicated DNA (see black dashed lines noting “after bypassing” in **Figure 7A** and **B**). Fortunately for cells, SMC loading is strongly biased to the *parS* sites that are closest to the replication origin, because *parS* sites are frequently very close to the replication origin ^29^, the most origin-proximal *parS* sites have higher binding affinity for ParB ^53^, and the origin region has the highest copy number in the chromosome (**Figure 1B**, MFA). Moreover, the rates of movement of the SMC complex and the replisome are closely matched and even in different species: for example, in *B. subtilis* growing at 42°C, replisomes move at ∼66 kb/min and SMC complex at ∼71 kb/min; in *Caulobacter crescentus*, replisomes move at ∼21 kb/min and SMC complex at ∼16-19 kb/min ^26, 54^. This close match of translocation rates of the two machines and the origin-biased SMC loading mean that in wild-type cells, the vast majority SMC complexes are “chasing” the replisome, and only occasionally catching up with them (**Figure 7C**). In this scenario, SMC bypassing the replisome would not result in unwanted chromosomal tethers or chromosome segregation issues (**Figure 7C**, after bypassing).

### SMC-replisome collisions in eukaryotes

In eukaryotes, SMC cohesins establish sister chromatid cohesion during DNA replication, thus the engagement between cohesin and the replisome has fundamental implications for DNA replication and segregation. Several recent *in vitro* single-molecule studies examined the outcomes of cohesin-replisome collisions. Using *Xenopus* egg extracts and a naked DNA substrate, it has been found that when colliding with the replisome head-on, cohesin is frequently blocked and pushed by the replisome to the converging point of replication forks. But at some frequency, cohesin unloads from the replisome, and at a lower frequency, cohesin bypasses the replisome ^55^. A separate study using *Xenopus* egg extract and a recent report using budding yeast cohesin both showed cohesion-replisome collision outcomes similar to the study mentioned above, although at different frequencies for pushing, unloading, and bypassing ^56, 57^. Besides these *in vitro* experiments, *in vivo* studies on the rules of engagement between cohesins and replisomes are lacking. Strikingly, our *in vivo* findings for SMC-replisome collision in bacteria mirror the *in vitro* observations for cohesin-replisome collision, establishing that the single-molecule results reflect what is happening inside cells. Moreover, our study provides interaction rules for replisome-SMC collisions that occur in the presence of all the other DNA transactions *in vivo*.

### Replisome movement is unaffected by SMC

Although SMC movement is restricted by the replisome, our experiments show that regardless of the location and the timing of SMC loading, the progression of DNA replication is unaffected (**Figures 1C, 2A, 6AB**). Single-molecule experiments also showed that replisomes were rarely affected by cohesins ^55, 57^. These findings highlight the mechanical prowess of the replisomes to clear or bypass protein roadblocks ^58^ ^59^, which is essential for DNA replication and genome integrity. A future challenge is to understand the molecular mechanism for SMC unloading and bypassing at the replisome, for instance, to identify the replisome component(s) responsible for SMC unloading, and to understand whether the ring-shaped SMC complex open when it bypasses the replisome. Further biochemical, structural, and single-molecule experiments are needed to address these questions.

In summary, replisome-mediated DNA replication and SMC-mediated chromosome organization and segregation are essential for cell viability. The resolution of collisions between the replisome and SMC complexes ensures faithful chromosome replication and segregation. We show that SMC complexes exhibit blocking, unloading, and bypassing behaviors when encountering the replisome *in vivo*, similar to *in vitro* results obtained for eukaryotic cohesin-replisome engagements. This convergence suggests that the interplay between SMC complexes and the replisome may represent a conserved mechanism for coordinating DNA replication and chromosome organization across diverse organisms.

## Acknowledgements

We thank the Wang lab for support and stimulating discussions, Leonid Mirny for computing resources, Xheni Karaboja for technical assistance, Alan Grossman for SMC antibodies, and Indiana University Center for Genomics and Bioinformatics high throughput sequencing. Support for this work comes from National Institutes of Health R01GM141242, R01GM143182, and R01AI172822 (X.W.). This research is a contribution of the GEMS Biology Integration Institute, funded by the National Science Foundation DBI Biology Integration Institutes Program, Award #2022049 (X.W.).

---

**Figure S1.**
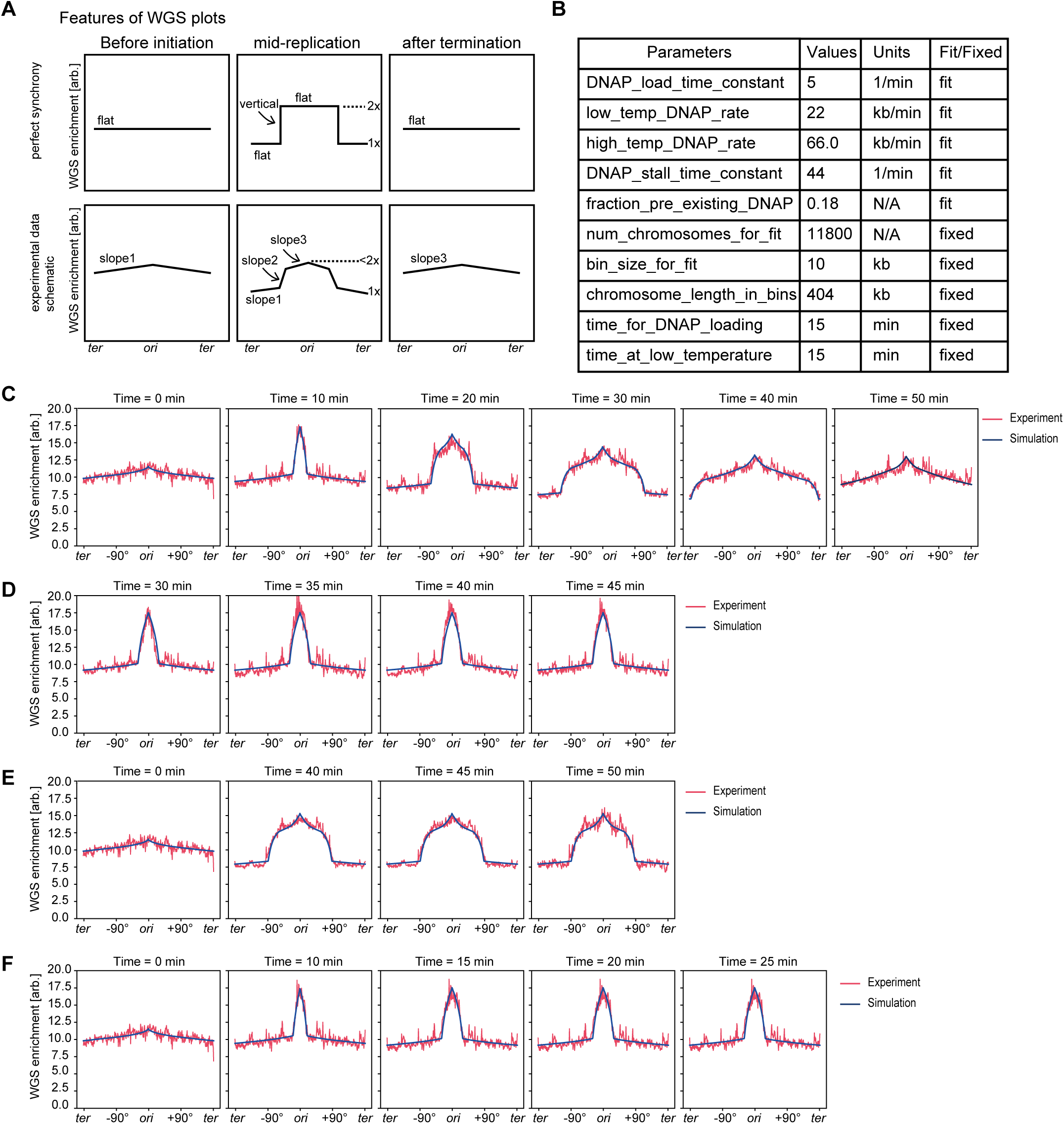
Determining replisome dynamics by simulating experimental MFA data. Related to Figure 1. **(A)** Illustrations depicting features of perfect replication synchrony (top) and the MFA plots in this study. **(B)** Parameters that produced the best fit for MFA plots used in **Figure 1C** (see the supplemental methods under “calibration of replisome dynamics model”), which are used for **C-F** below. **(C)** Comparison of simulated replication profiles (blue curves) with experimental MFA plots (red curves) for the strain in **Figure 1C**. **(D-F)** Comparison of simulated replication profiles (blue curves) with experimental MFA plots (red curves) for experiments presented in **Figure 4D**, **4E** and **4F**, respectively.

**Figure S2.**
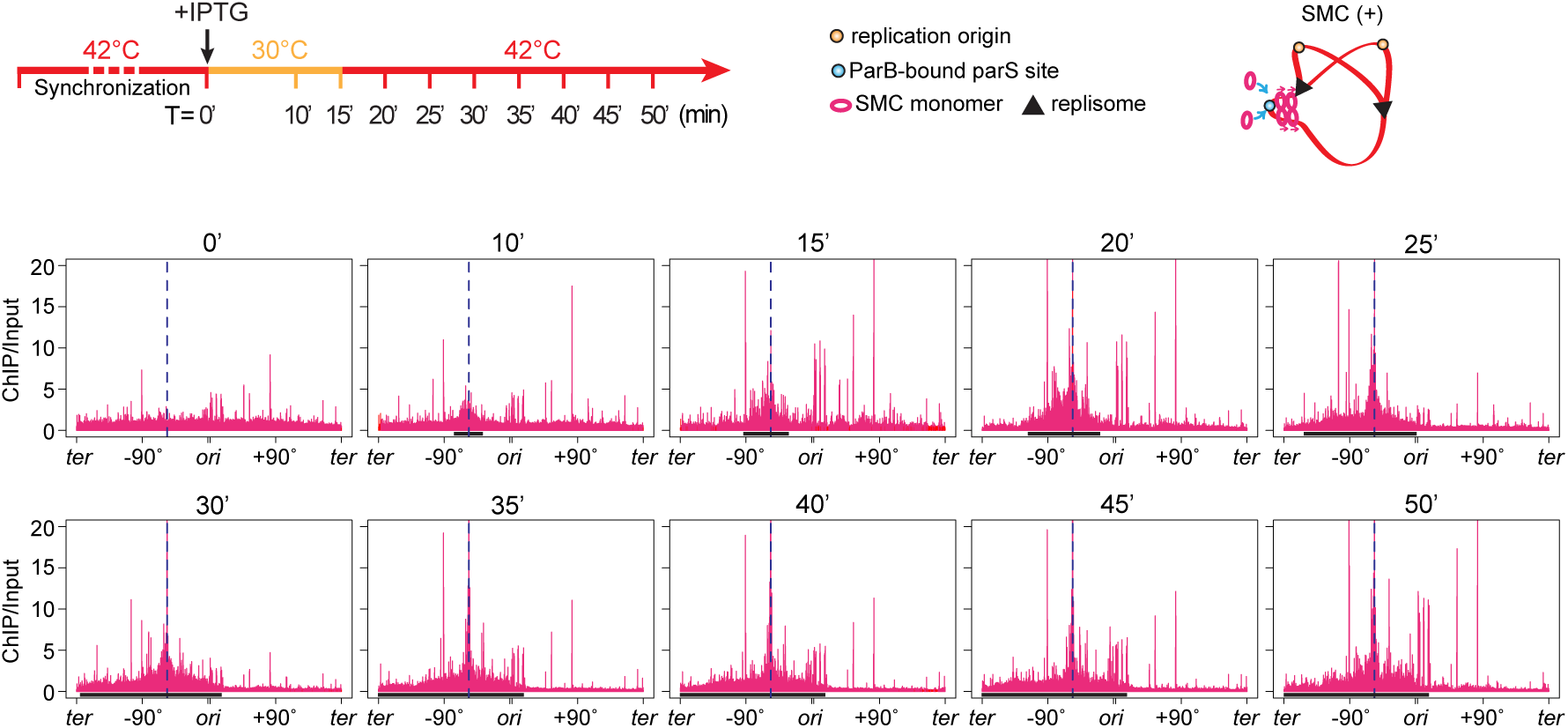
SMC enrichment upon ParB expression in a strain containing *parS* at -59° (BWX5297). Related to **Figure 1**. SMC ChIP-seq enrichment profiles of the samples used in **Figure 1C**. ChIP enrichments (ChIP/input) were plotted in 1-kb bins. The *parS* site is indicated by a blue dashed line. SMC enrichment zones are indicated as black bars in this figure, but as magenta bars in MFA plots in **Figure 1C**.

**Figure S3.**
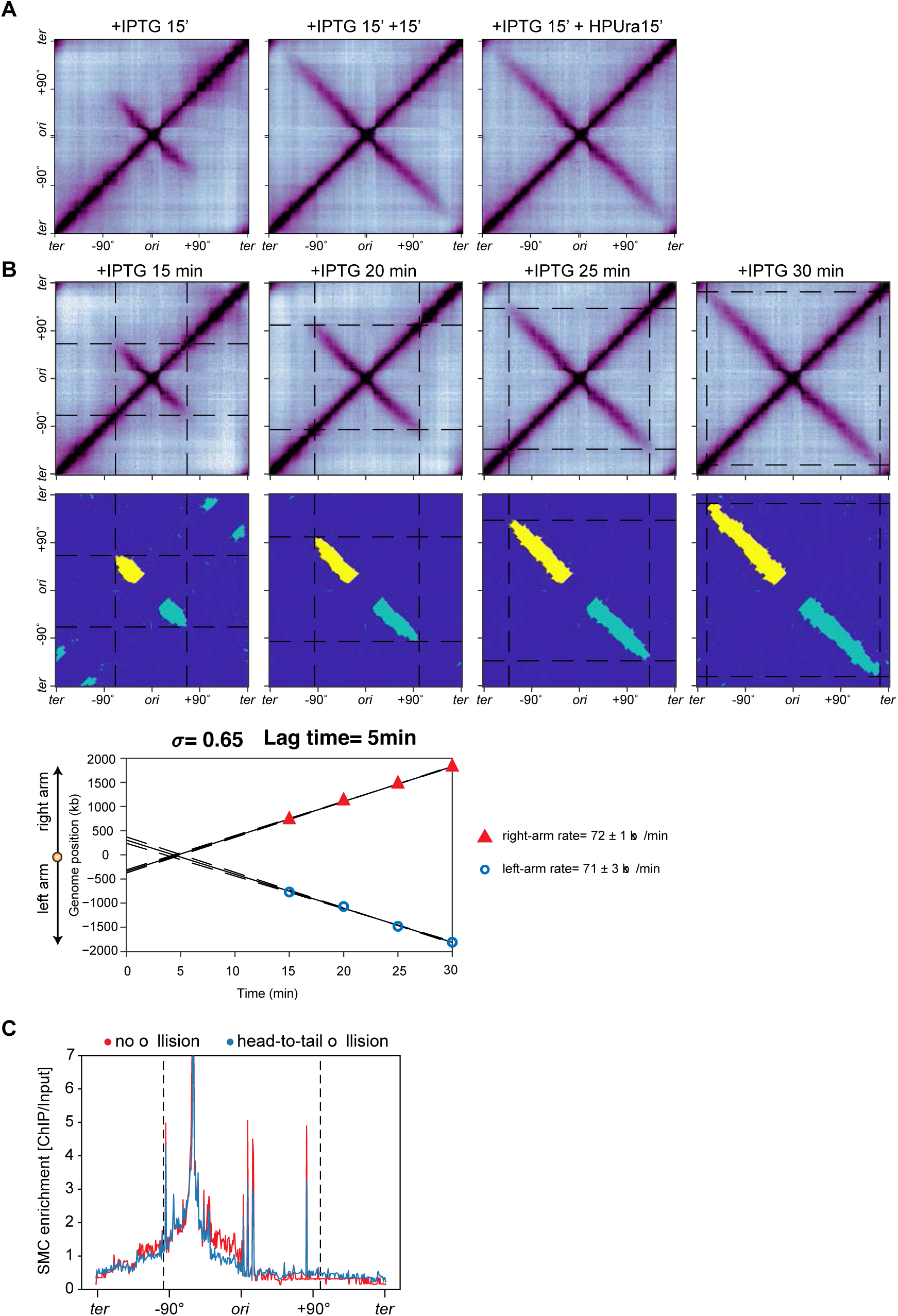
HPUra does not affect SMC translocation rate. Related to **Figure 3** and 4. **(A)** Hi-C maps of a strain containing a single *parS* site at -1° and IPTG-inducible *parB* (BWX4310). Cells were arrested in G1. IPTG were added for 15 min to induce SMC loading, then cells were treated with or without HPUra for another 15 min. Without ongoing replication, SMC-mediated DNA juxtaposition was not affected by HPUra. **(B)** The DNA juxtaposition rate at 42°C without SMC-replisome collisions was determined as described previously ^1^. 0.65x standard deviation (s) above the averaged Hi-C contact score was used as the threshold. Top: time-course Hi-C maps shown in **Figure 4A**. Middle: binary maps showing points with Hi-C contact scores higher than the threshold. Bottom: quantified DNA juxtaposition at indicated time points. The SMC extrusion rate was calculated from the slope of the lines in the plot. **(C)** A comparison of SMC ChIP enrichment profiles shown in **Figure 4C** (red) and **Figure 4D** (blue) for the +IPTG 25 min time point. ChIP enrichments (ChIP/input) were plotted in 10-kb bins. The positions of replication forks are indicated by black dashed lines.

**Figure S4.**
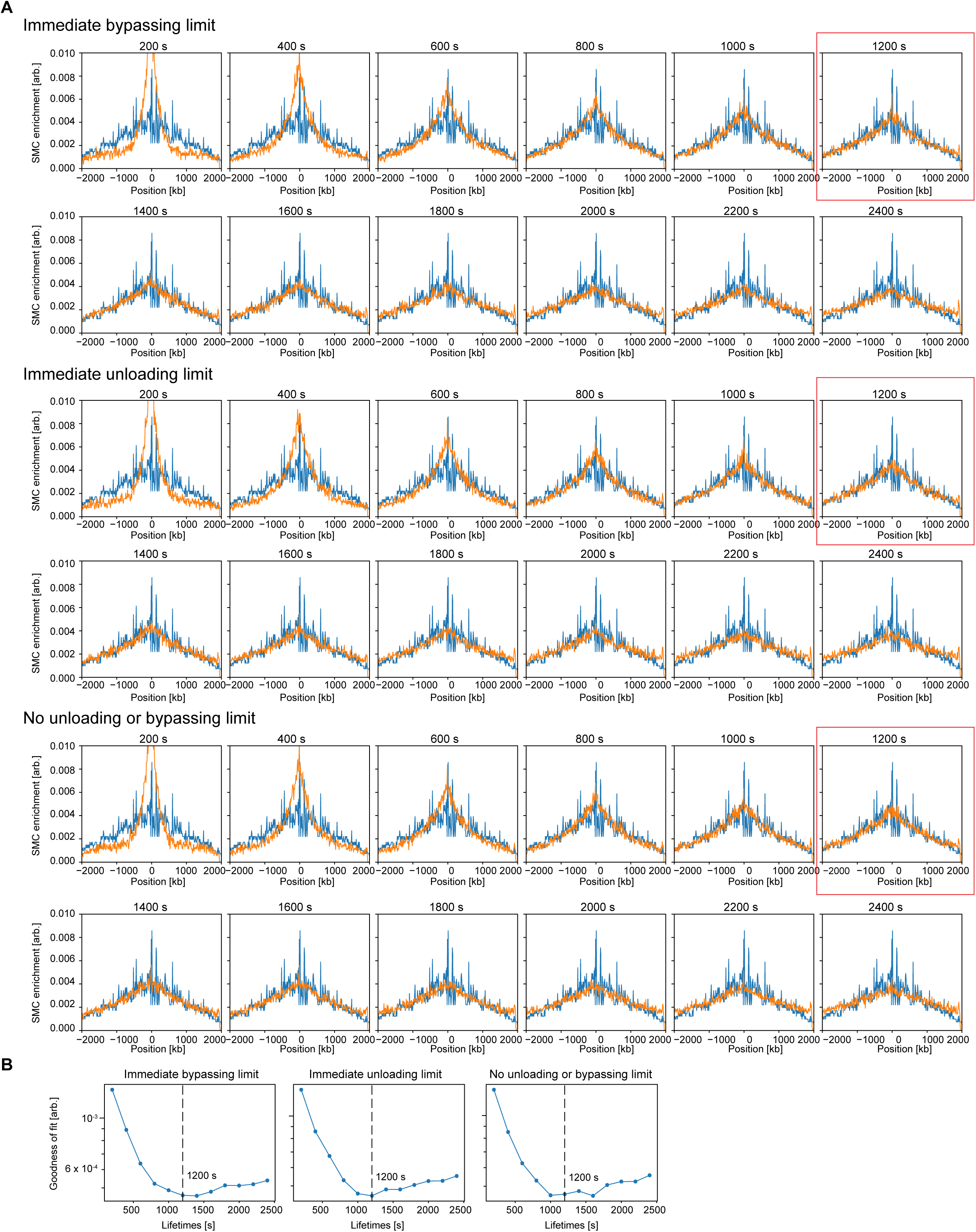
Calibration of SMC spontaneous disassociation rate at 42°C. Related to Figure 5. **(A)** SMC occupancy (orange curves) was simulated and compared with experimental ChIP-seq result (blue curves) that was obtained using G1-arrested cells growing at 42°C. This strain contained a single *parS* site at -1° and IPTG-inducible *sirA* (BWX4504), which inhibited replication initiation after 1 h in the presence of 1 mM IPTG. Since a small portion of cells had pre-existing replisomes as indicated in MFA results, we considered three scenarios of SMC-replisome interaction upon the encounter: 1) SMC bypasses the replisome; 2) SMC unloads from the chromosome; 3) SMC is blocked by the replisome. Simulated SMC occupancy was obtained by varying the SMC spontaneous dissociation rate from 1/200 s^-1^ to 1/1200 s^-1^. See details in the supplemental methods under “calibration of spontaneous dissociation rate of SMC complexes”. **(B)** Goodness-of-fit analysis to compare the deviations between simulated SMC enrichments (show in **A**) and the experimental ChIP data. The best fit is the one with the minimum score. Combined with visual inspection of SMC enrichment profiles, we chose 1/1200 s^-1^ as the spontaneous dissociation rate in all experiments.

**Figure S5.**
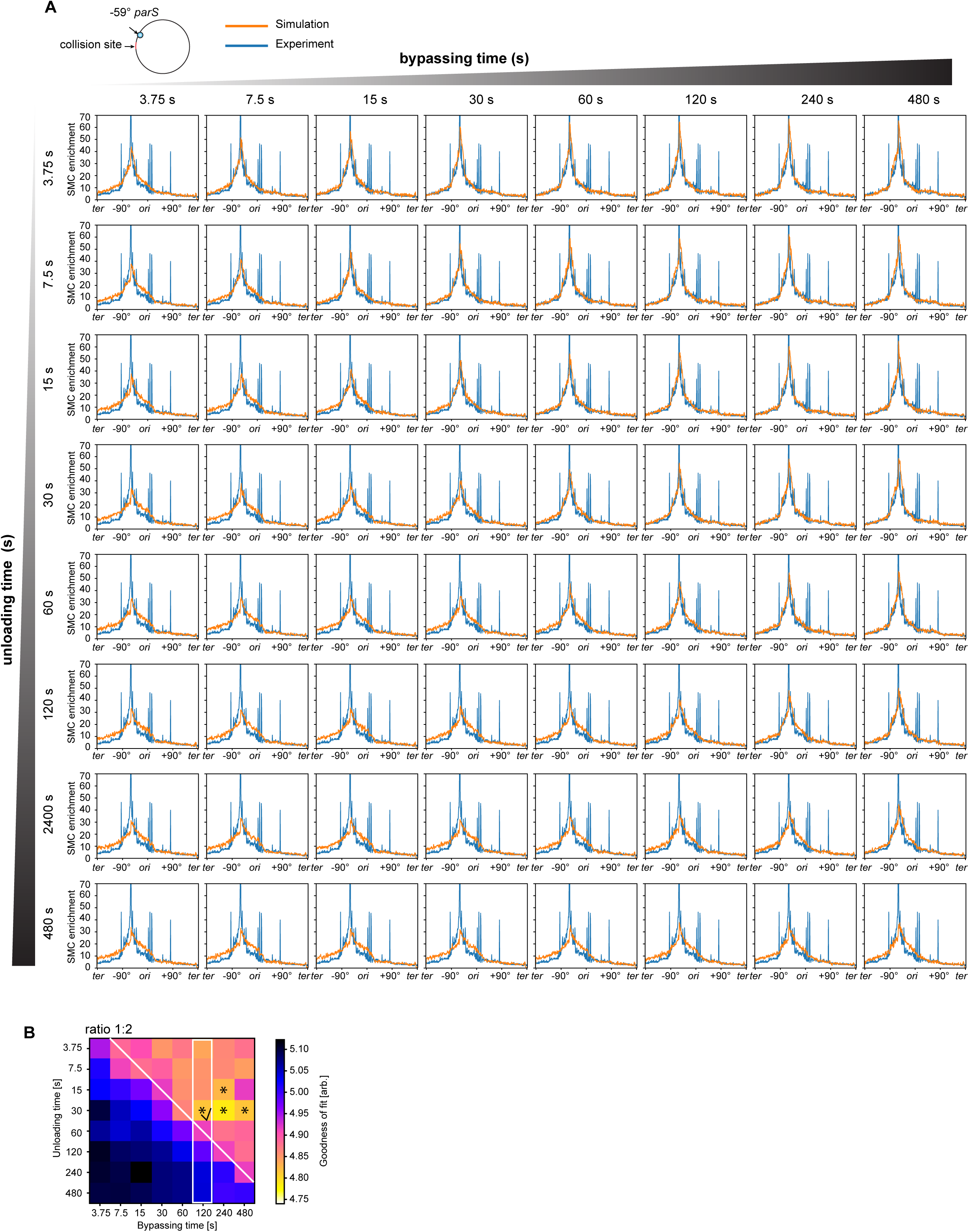
Sweeping of bypassing rates and unloading rates in the one-sided head-to-tail collision. Related to Figure 5A-E. **(A)** SMC occupancy (orange curves) was simulated by varying bypassing time and unloading time, and compared with experimental ChIP-seq result (**Figure 4D**, +IPTG 25 min time point; **Figure 5A**) (blue curves). See details the supplemental methods under “parameter sweeps for interaction rules between SMC and the stalled replisome”. **(B)** Goodness-of-fit heatmap generated from simulations in (**A**). The white diagonal line indicates the 1:2 ratio of unloading time to bypassing time. The four combinations that produced the best fit are highlighted with stars. Since the experimental Hi-C analysis estimated the bypassing time to be about two minutes for this one-sided head-to-tail collision (white rectangle), the best combination in the range is unloading at 30 s and bypassing at 120 s (black check mark).

**Figure S6.**
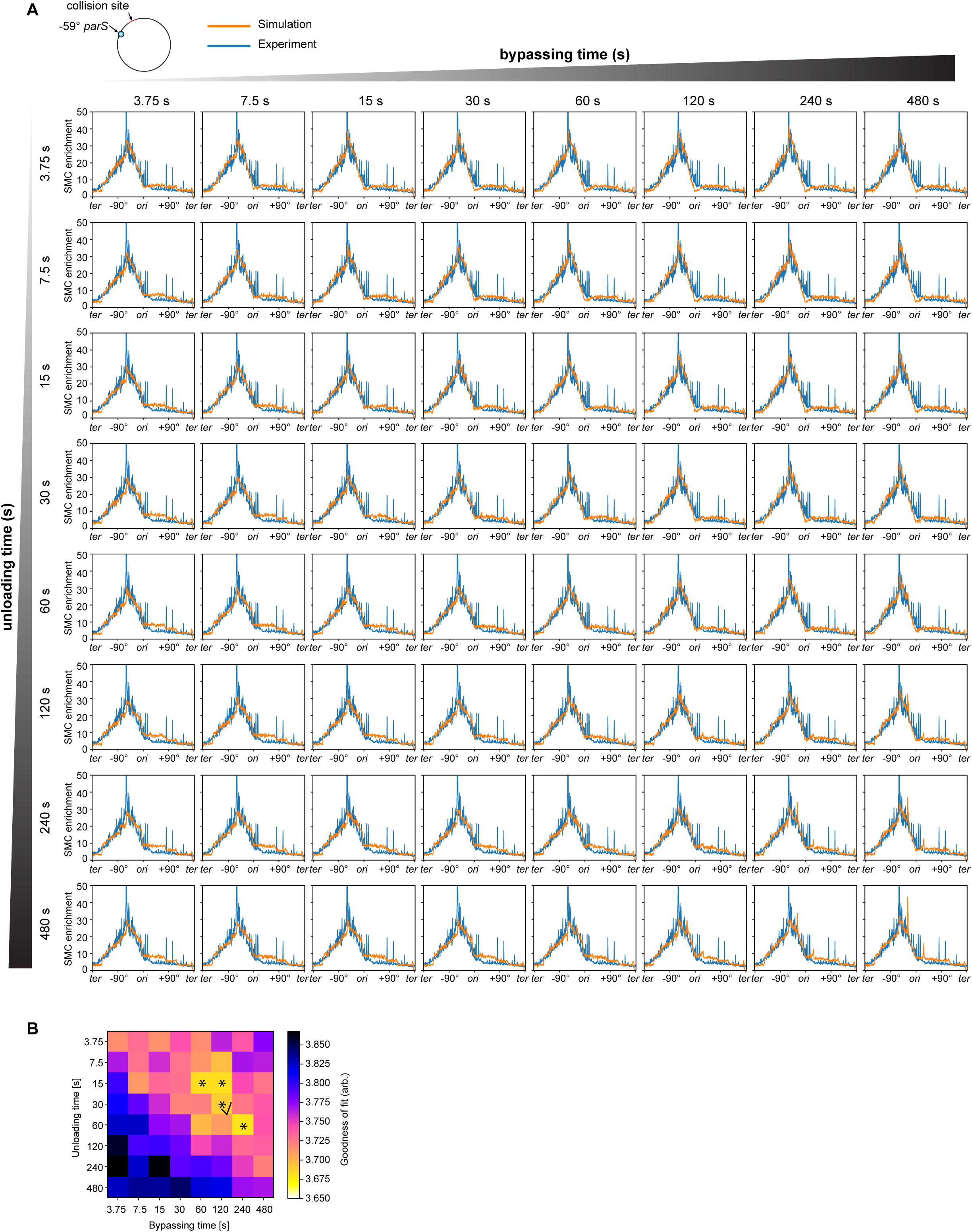
Sweeping of bypassing rates and unloading rates in the head-on collision. Related to Figure 5F-J. **(A)** SMC occupancy (orange curves) was simulated by varying bypassing time and unloading time, and compared with experimental ChIP-seq result (**Figure 4E**, +IPTG 25 min time point; **Figure 5F**) (blue curves). See details in the supplemental methods under “parameter sweeps for interaction rules between SMC and the stalled replisome”. **(B)** Goodness-of-fit heatmap generated from simulations in (**A**). The four combinations that produced the best fit are highlighted with stars. The same combination used for head-to-tail collision in **Figure S5** (unloading=30 s and bypassing=120 s) is indicated by a black check mark.

**Figure S7.**
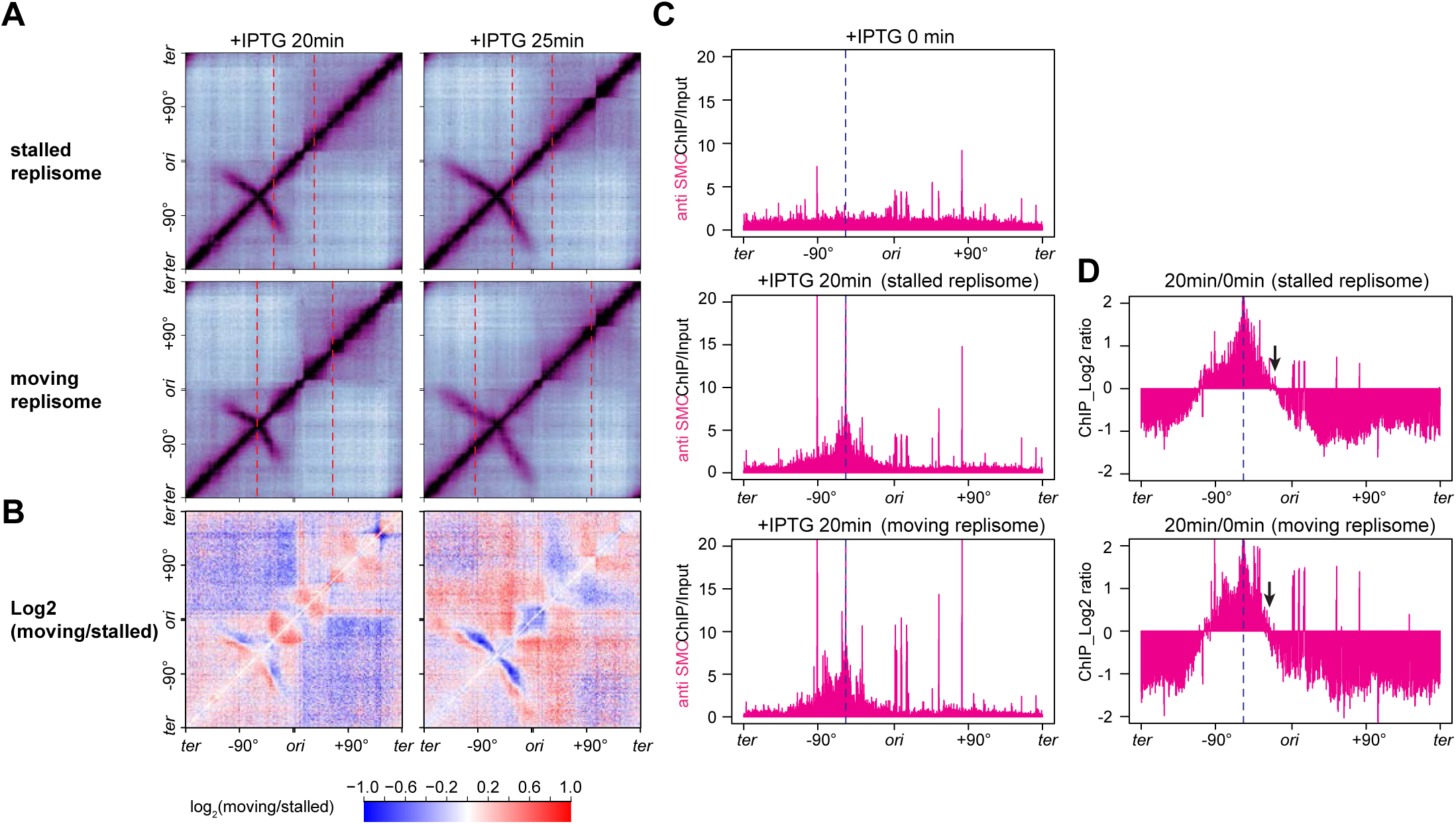
Moving replisomes have a stronger effect on SMC than stalled replisomes. Related to Figure 6. **(A)** Hi-C maps obtained from SMC collisions with stalled replisomes (top, **Figure 4E**, time points 20 min and 25 min) and from SMC collisions with moving replisomes (bottom, **Figure 6A**, time points + IPTG 20 min and +IPTG 25 min). Red dashed lines indicate the position of replisomes determined by MFA plots. **(B)** Log2 ratio of Hi-C matrices between “moving replisome” and “stalled replisome” shown in (**A**). Red pixels designate increased interactions while blue pixels indicate decreased interactions. **(C)** SMC ChIP enrichment before IPTG induction (top), or after 20-min IPTG induction in **Figure 4E** which had stalled replisomes, or 20-min IPTG induction in **Figure 6A** which had moving replisomes. Blue dashed lines indicate the position of the *parS* site. **(D)** The ratio of ChIP enrichment at 20 min relative to 0 min were plotted in log2 scale in 1-kb bins to track SMC translocation after induction. The top graph used cells containing stalled replisomes and the bottom graph used cells containing moving replisomes. Black arrows indicate the endpoints of SMC enrichment toward the replication origin.

**Table S1.**
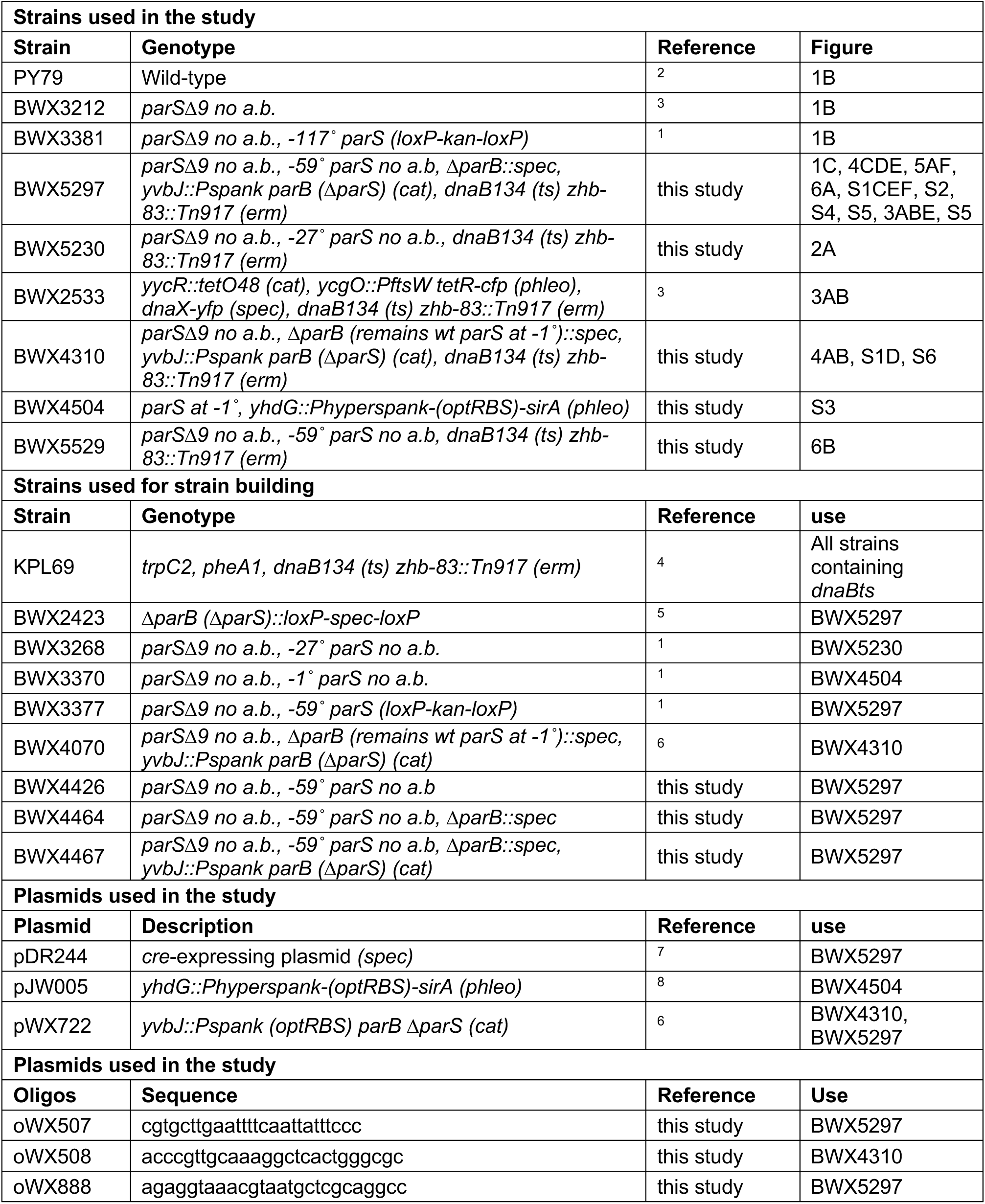
Bacterial strains and plasmid used in this study.

**Table S2.** Next-Generation-Sequencing data used in this study.

| **Sample name** | **Figure** | **Reference** |
| --- | --- | --- |
| WGS_PY79_exp | 1B | This study |
| WGS_BWX3212_exp | 1B | This study |
| WGS_BWX3381_exp | 1B | This study |
| WGS_BWX5297_T0 | 1C, 4C, 6A, S2, S7B | This study |
| WGS_BWX5297_30C_IPTG10min_T10 | 1C, 6A, S2 | This study |
| WGS_BWX5297_30C_IPTG15min_T15 | 1C, 4E, 6A, S2 | This study |
| WGS_BWX5297_42C_IPTG20min_T20 | 1C, 6A, S2, S7B | This study |
| WGS_BWX5297_42C_IPTG25min_T25 | 1C, 6A, S2 | This study |
| WGS_BWX5297_42C_IPTG30min_T30 | 1C, 6A, S2 | This study |
| WGS_BWX5297_42C_IPTG35min_T35 | 1C, 6A, S2 | This study |
| WGS_BWX5297_42C_IPTG40min_T40 | 1C, 6A, S2 | This study |
| WGS_BWX5297_42C_IPTG45min_T45 | 1C, 6A, S2 | This study |
| WGS_BWX5297_42C_IPTG50min_T50 | 1C, 6A, S2 | This study |
| WGS_BWX5297_30C_T10 | 1C | This study |
| WGS_BWX5297_30C_T15 | 1C | This study |
| WGS_BWX5297_42C_T20 | 1C | This study |
| WGS_BWX5297_42C_T25 | 1C | This study |
| WGS_BWX5297_42C_T30 | 1C | This study |
| WGS_BWX5297_42C_T35 | 1C | This study |
| WGS_BWX5297_42C_T40 | 1C | This study |
| WGS_BWX5297_42C_T45 | 1C | This study |
| WGS_BWX5297_42C_T50 | 1C | This study |
| WGS_BWX5230_T0 | 2A | This study |
| WGS_BWX5230_30C_T10 | 2A | This study |
| WGS_BWX5230_30C_T15 | 2A | This study |
| WGS_BWX5230_42C_T20 | 2A | This study |
| WGS_BWX5230_42C_T25 | 2A | This study |
| WGS_BWX5230_42C_T30 | 2A | This study |
| WGS_BWX5230_42C_T35 | 2A | This study |
| WGS_BWX5230_42C_T40 | 2A | This study |
| WGS_BWX4310_exp | 3A | This study |
| WGS_BWX4310_T0 | 3A | This study |
| WGS_BWX4310_30C_T15 | 3A | This study |
| WGS_BWX4310_42C_IPTG15min_T30 | 3A, 4A | This study |
| WGS_BWX4310_42C_IPTG30min_T45 | 3A, 4A | This study |
| WGS_BWX4310_42C_HPUraIPTG15min_T30 | 3A, 4B | This study |
| WGS_BWX4310_42C_HPUraIPTG30min_T45 | 3A, 4B | This study |
| WGS_BWX4310_42C_IPTG20min_T35 | 4A | This study |
| WGS_BWX4310_42C_IPTG25min_T40 | 4A | This study |

| Sample name | Figure | Reference |
| --- | --- | --- |
| WGS_BWX4310_42C_HPUraIPTG20min_T35 | 4B | This study |
| WGS_BWX4310_42C_HPUraIPTG25min_T40 | 4B | This study |
| WGS_BWX5297_42C_noreplisome_IPTG15min_T15 | 4C | This study |
| WGS_BWX5297_42C_noreplisome_IPTG20min_T20 | 4C | This study |
| WGS_BWX5297_42C_noreplisome_IPTG25min_T25 | 4C | This study |
| WGS_BWX5297_42C_HPUraIPTG15min_T40 | 4D | This study |
| WGS_BWX5297_42C_HPUraIPTG20min_T45 | 4D | This study |
| WGS_BWX5297_42C_HPUraIPTG25min_T50 | 4D | This study |
| WGS_BWX5297_42C_IPTG20min_HPUra5min_T20 | 4E, S7B | This study |
| WGS_BWX5297_42C_IPTG25min_HPUra10min_T25 | 4E | This study |
| WGS_BWX5529_T0 | 6B | This study |
| WGS_BWX5529_30C_T10 | 6B | This study |
| WGS_BWX5529_30C_T15 | 6B | This study |
| WGS_BWX5529_42C_T20 | 6B | This study |
| WGS_BWX5529_42C_T25 | 6B | This study |
| WGS_BWX5529_42C_T30 | 6B | This study |
| WGS_BWX5529_42C_T35 | 6B | This study |
| WGS_BWX5529_42C_T40 | 6B | This study |
| WGS_BWX4504_42C_IPTG1h | S4 | This study |
| ChIP_anti_SMC_BWX5297_T0 | S2, S7B | This study |
| ChIP_anti_SMC_BWX5297_30C_IPTG10min_T10 | S2 | This study |
| ChIP_anti_SMC_BWX5297_30C_IPTG15min_T15 | S2 | This study |
| ChIP_anti_SMC_BWX5297_42C_IPTG20min_T20 | S2, S7B | This study |
| ChIP_anti_SMC_BWX5297_42C_IPTG25min_T25 | S2 | This study |
| ChIP_anti_SMC_BWX5297_42C_IPTG30min_T30 | S2 | This study |
| ChIP_anti_SMC_BWX5297_42C_IPTG35min_T35 | S2 | This study |
| ChIP_anti_SMC_BWX5297_42C_IPTG40min_T40 | S2 | This study |
| ChIP_anti_SMC_BWX5297_42C_IPTG45min_T45 | S2 | This study |
| ChIP_anti_SMC_BWX5297_42C_IPTG50min_T50 | S2 | This study |
| ChIP_anti_SMC_BWX5297_42C_noreplisome_IPTG25min_T25 | S3 | This study |
| ChIP_anti_SMC_BWX5297_42C_HPUraIPTG25min_T50 | 5A, S5 | This study |
| ChIP_anti_SMC_BWX5297_42C_IPTG25min_HPUra10min_T25 | 5B, S6 | This study |
| ChIP_anti_SMC_BWX4504_42C_IPTG1h | S4 | This study |
| ChIP_anti_SMC_BWX5297_42C_IPTG20min_HPUra5min_T20 | S7B | This study |
| HiC_BWX5230_T0 | 2A | This study |
| HiC_BWX5230_30C_T10 | 2A | This study |
| HiC_BWX5230_30C_T15 | 2A | This study |
| HiC_BWX5230_42C_T20 | 2A | This study |
| HiC_BWX5230_42C_T25 | 2A | This study |
| HiC_BWX5230_42C_T30 | 2A | This study |
| HiC_BWX5230_42C_T35 | 2A | This study |
| HiC_BWX5230_42C_T40 | 2A | This study |
| HiC_BWX4310_42C_IPTG15min_T30 | 4A, S3B | This study |
| HiC_BWX4310_42C_IPTG20min_T35 | 4A, S3B | This study |
| HiC_BWX4310_42C_IPTG25min_T40 | 4A, S3B | This study |
| HiC_BWX4310_42C_IPTG30min_T45 | 4A, S3B | This study |
| HiC_BWX4310_42C_HPUraIPTG15min_T30 | 4B | This study |
| HiC_BWX4310_42C_HPUraIPTG20min_T35 | 4B | This study |
| HiC_BWX4310_42C_HPUraIPTG25min_T40 | 4B | This study |
| HiC_BWX4310_42C_HPUraIPTG30min_T45 | 4B | This study |
| HiC_BWX5297_T0 | 4CDE, 6A | This study |
| HiC_BWX5297_42C_noreplisome_IPTG15min_T15 | 4C | This study |
| HiC_BWX5297_42C_noreplisome_IPTG20min_T20 | 4C | This study |
| HiC_BWX5297_42C_noreplisome_IPTG25min_T25 | 4C | This study |
| HiC_BWX5297_42C_HPUraIPTG15min_T40 | 4D | This study |
| HiC_BWX5297_42C_HPUraIPTG20min_T45 | 4D | This study |
| HiC_BWX5297_42C_HPUraIPTG25min_T50 | 4D | This study |
| HiC_BWX5297_30C_IPTG15min_T15 | 4E, 6A | This study |
| HiC_BWX5297_42C_IPTG20min_HPUra5min_T20 | 4E | This study |
| HiC_BWX5297_42C_IPTG25min_HPUra10min_T25 | 4E | This study |
| HiC_BWX5297_30C_IPTG10min_T10 | 6A | This study |
| HiC_BWX5297_42C_IPTG20min_T20 | 6A | This study |
| HiC_BWX5297_42C_IPTG25min_T25 | 6A | This study |
| HiC_BWX5297_42C_IPTG30min_T30 | 6A | This study |
| HiC_BWX5297_42C_IPTG35min_T35 | 6A | This study |
| HiC_BWX5297_42C_IPTG40min_T40 | 6A | This study |
| HiC_BWX5529_T0 | 6B | This study |
| HiC_BWX5529_30C_T10 | 6B | This study |
| HiC_BWX5529_30C_T15 | 6B | This study |
| HiC_BWX5529_42C_T20 | 6B | This study |
| HiC_BWX5529_42C_T25 | 6B | This study |
| HiC_BWX5529_42C_T30 | 6B | This study |
| HiC_BWX5529_42C_T35 | 6B | This study |
| HiC_BWX5529_42C_T40 | 6B | This study |
| Hi-C_BWX4310_42C_IPTG15min | S3 | This study |
| Hi-C_BWX4310_42C_IPTG15min_HPUra15min | S3 | This study |
| Hi-C_BWX4310_42C_IPTG30min | S3 | This study |

## Supplemental Methods

### General Methods

*Bacillus subtilis* strains were derived from the prototrophic strain PY79 ^2^. Cells were grown in defined rich Casein Hydrolysate (CH) medium at 22°C, 30°C or 42°C as indicated. ParB production was induced by the addition of 1 mM IPTG (Dot Scientific, DS102125). To arrest replisomes movement, 162 µM HPUra ^9, 10, 11^ was used. Lists of strains, plasmids and oligonucleotides can be found in Table S1. List of Next-Generation-Sequencing samples can be found in Tables S2.

### Hi-C

The Hi-C procedure was carried out as previously described ^3, 12^. Specifically, cells grown at the desired condition were crosslinked with 3% formaldehyde at room temperature for 30 min then quenched with 125 mM glycine. Cells were lysed using Ready-Lyse Lysozyme (Epicentre, R1802M) and treated by 0.5% SDS. Solubilized chromatin was digested with HindIII for two hours at 37°C. The digested ends were filled in with Klenow and Biotin-14-dATP, dGTP, dCTP, dTTP. The products were ligated with T4 DNA ligase at 16°C for about 20 h. Crosslinks were reversed at 65°C for about 20 h in the presence of EDTA, proteinase K and 0.5% SDS. The DNA was then extracted twice with phenol/chloroform/isoamylalcohol (25:24:1) (PCI), precipitated with ethanol, and resuspended in 20 µl of 0.1X TE buffer. Biotin from non-ligated ends was removed using T4 polymerase (4 h at 20°C) followed by extraction with PCI. The DNA was then sheared by sonication for 12 min with 20% amplitude using a Qsonica Q800R2 water bath sonicator. The sheared DNA was used for library preparation with the NEBNext UltraII kit (E7645).

Biotinylated DNA fragments were purified using 5 µl streptavidin beads. DNA-bound beads were used for PCR in a 50 µl reaction for 14 cycles. PCR products were purified using Ampure beads (Beckman, A63881) and sequenced at the Indiana University Center for Genomics and Bioinformatics using NextSeq500. Paired-end sequencing reads were mapped to the genome of *B. subtilis* PY79 (NCBI Reference Sequence NC_022898.1) using the same pipeline described previously ^3^. The genome was divided into 10-kb bins. Subsequent analysis and visualization were done using R scripts.

### ChIP-seq

Chromatin immunoprecipitation (ChIP) was performed as described previously ^3^. Briefly, cells were crosslinked using 3% formaldehyde for 30 min at room temperature and then quenched using 125 mM glycine, washed using PBS, and lysed using lysozyme. Crosslinked chromatin was sheared to an average size of 250 bp by sonication using Qsonica Q800R2 water bath sonicator. The lysate was precleared using Protein A magnetic beads (GE Healthcare/cytiva 28951378) and was then incubated with anti-SMC antibody ^13^ overnight at 4°C. Next day, the lysate was incubated with Protein A magnetic beads for one hour at 4°C. After washes and elution, the immunoprecipitate was incubated at 65°C overnight to reverse the crosslinks. The DNA was further treated with RNaseA, Proteinase K, extracted with PCI, resuspended in 100 µl EB and used for library preparation with the NEBNext UltraII kit (E7645). The library was sequenced at the Indiana University Center for Genomics and Bioinformatics using NextSeq500. The sequencing reads were mapped to the genome of *B. subtilis* PY79 (NCBI Reference Sequence NC_022898.1) using CLC Genomics Workbench (CLC Bio, QIAGEN). Sequencing reads from each ChIP and input sample were normalized by the total number of reads. The ChIP enrichment (ChIP/Input) was plotted and analyzed using R scripts. For comparisons of experimental ChIP enrichment profiles to the ones generated from simulations, the experimental data were filtered and plotted as described in the method section “*Filtering of ChIP-seq and MFA data for goodness-of-fit calculations and visual comparison*”.

### Whole Genome Sequencing (WGS)

Cells were grown and collected at indicated time points under the desired conditions. Collected cells were crosslinked with 3% formaldehyde at room temperature for 30 min then quenched with 125 mM glycine. Cells were lysed using Ready-Lyse Lysozyme (Epicentre, R1802M) and treated by 0.5% SDS. Cell lysate was incubated at 65°C overnight to reverse the crosslinks. The DNA was further treated with RNaseA, Proteinase K, extracted with PCI, resuspended in 200 µl of 0.1X TE buffer. The extracted DNA was sonicated using a Qsonica Q800R2 water bath sonicator, prepared using the NEBNext UltraII kit (E7645), and sequenced at the Indiana University Center for Genomics and Bioinformatics using NextSeq500. The reads were mapped to the genome of *B. subtilis* PY79 (NCBI Reference Sequence NC_022898.1) using CLC Genomics Workbench (CLC Bio, QIAGEN). The mapped reads were normalized by the total number of reads. Plotting and analysis were performed using R scripts. For comparisons of experimental WGS profiles to the ones generated from simulations, the experimental data were filtered and plotted as described in the method section “*Filtering of ChIP-seq and MFA data for goodness of fit calculations and visual comparison*”.

### Fluorescence microscopy analysis

Fluorescence microscopy was performed on a Nikon Ti2E microscope equipped with Plan Apo 100x/1.4NA phase contrast oil objective and an sCMOS camera. 0.2 OD unit of cells were collected for the imaging. Membranes were stained with FM4-64 (Molecular Probes) at 3 μg/ml. DNA was stained with DAPI (Molecular Probes) at 2 μg/ml.

### Strain construction

#### -59° parS, ΔparB, Pspank parB without parS, dnaB (ts) (BWX5297)

This strain was constructed in four steps: 1) The *loxP-kan-loxP* cassette in BWX3377 ^1^ was removed using a *cre*-expressing plasmid pDR244 ^7^, resulting in BWX4426; 2) A PCR product of Δ*parB* clean deletion was amplified using oWX507 and oWX888 from strain BWX2423 ^5^ and transformed into strain BWX4426, resulting in BWX4464; 3) pWX722 (*Pspank parB without parS*) ^6^ was transformed into BWX4464, resulting in BWX4467; 4) The genomic DNA of *dnaB134 (ts) zhb-83::Tn917* ^4^ was transformed into BWX4467.

#### -59° parS, dnaB (ts) (BWX5529)

This strain was constructed by transforming the genomic DNA of the *dnaB134 (ts) zhb-83::Tn917* strain ^4^ into BWX4426 described above.

#### -27° parS, dnaB (ts) (BWX5230)

This strain was constructed by transforming the genomic DNA of the *dnaB134 (ts) zhb-83::Tn917* strain ^4^ into BWX3268 (*parSΔ9 no a.b., -27° parS no a.b.*) ^1^.

#### -1° parS, ΔparB, Pspank parB without parS, dnaB (ts) (BWX4310)

This strain was constructed by transforming the genomic DNA of the *dnaB134 (ts) zhb-83::Tn917* strain ^4^ into BWX4070 ^6^.

#### -1° parS, Phyperspank sirA (BWX4504)

This strain was constructed by transforming pJW005 ^8^ into BWX3370 ^1^.

### Calibration of replisome dynamics model

We first simulated replisome dynamics in a population of synchronously growing cells. Our objective was to reproduce the shape of the marker-frequency analysis (MFA) as determined by whole-genome sequencing (WGS) to define the replisome distribution on the genome using the fewest parameters possible.

We used the MFA data from the *dnaB (ts)* strain (BWX5297) in **Figure 1C**. We performed simulations to vary several parameters on replisome behaviors: 1) the loading rate of DNA polymerase (DNAP) to the origin (*DNAP_load_time*); 2) the speed of replication forks at 30°C (*low_temp_DNAP_rate*); 3) the speed of replication forks at 42°C (*high_temp_DNAP_rate*); 4) the time for spontaneous DNAP stalling (*DNAP_stall_time*); 5) the fraction of cells containing replisomes before initiation (*fraction_pre-existing DNAP*). See below for detailed explanations.

In the experiment in **Figure 1C**, cells were first synchronized at the G-1 stage by being grown at 42°C for 45 min (T=0 min). The overall “flat” shape of the curve in the MFA profile at this time point (**Figure 1C**, 0’) indicates that most of the cells in the synchronous population contained a single chromosome. However, the slight “hat shape” (i.e. ^) of the curve (**Figure S1A**, slope 1) indicates that a some cells had partially replicated chromosome prior to replication initiation, despite growing at 42°C. We refer to the replisomes in this subpopulation as pre-existing replisomes, heuristically modeled its fraction by the *fraction_pre_existing_DNAP* parameter, and distributed these replisomes using a wide quadratic distribution centered with its maximum at the origin.

For experimental data upon replication initiation, we found that the shapes of the MFA distribution were best matched by including a parameter, *DNAP_stall_time*, to account for spontaneous DNAP stalling. This parameter modeled the probability that a replisome prematurely stops, which could be caused by factors such as DNA damage or other temporary roadblocks. The spontaneous stalling time was modeled using an exponential distribution with the characteristic stalling rate governed by 1/*DNAP_stall_time*.

We combined all these elements into a function to compute the probability distribution of DNAP positions along the chromosome at given time points. Briefly, we uniformly sampled the loading times for each DNAP from an exponential distribution with average time *DNAP_load_time* to pre-determine the loading time of each replisome (DNAP). Additionally, we sampled *DNAP_stall_time* from an exponential distribution with average time to determine the spontaneous stalling time of each replisome (DNAP). We then paired up all possible combinations of sampled DNAP loading times and stalling times to generate a minimum number of sampled (simulated) replisomes.

Upon replication initiation, for simplicity, we assumed that replisomes progressed symmetrically to the left and right of the origin. Each replisome traveled away from the origin using the rate *low_temp_DNAP_rate* for the first 15 min when the cells were grown at 30°C, followed by *high_temp_DNAP_rate* when growing 42°C. The distance traveled by each replisome arm was computed by (*low_temp_DNAP_rate* * 10 min or 15 min and *high_temp_DNAP_rate* * time for time T-15 min). A fraction of pre-existing chromosomes was additionally distributed as described above.

This model for the distribution of replisome positions was then normalized and compared to experimental data. To fit the model parameters, we employed a least-squares optimization approach to collectively minimize the difference between the distributions of simulated replisome position and the experimental MFA data at the time points 0 min, 10 min, 20 min, 30 min, 40 min, 50 min. Both simulation and experimental MFA data were binned into 10 kb segments for a total of 404 bins for the whole genome.

We normalized both the experimental data and the simulated copy number by the sum of total counts, then multiplied the values by 4200. This was calculated by: 42 * 1e6/1e4 = 4200. The factor of 42 comes from the fact that the WGS read length is 42 bases long; the 1e6 is for reporting values in reads per million; finally, we divide the total by 1e4 since the data is median filtered with a window size of 10 kb. For the least-squares optimization, we used scipy1.4.1 (scipy.optimize.minimize). By sweeping the parameters above, we identified a set of best-fit values that reproduced the experimental MFA data in **Figure 1C** (also see **Figure S1B**, and **C**).

| Parameters | Values | Units | Fit/Fixed |
| --- | --- | --- | --- |
| DNAP_load_time_constant | 5 | 1/min | fit |
| low_temp_DNAP_rate | 22 | kb/min | fit |
| high_temp_DNAP_rate | 66.0 | kb/min | fit |
| DNAP_stall_time_constant | 44 | 1/min | fit |
| fraction_pre_existing_DNAP | 0.18 | N/A | fit |
| num_chromosomes_for_fit | 11800 | N/A | fixed |
| bin_size_for_fit | 10 | kb | fixed |
| chromosome_length_in_bins | 404 | kb | fixed |
| time_for_DNAP_loading | 15 | min | fixed |
| time_at_low_temperature | 15 | min | fixed |

Using the same fitting procedure and same parameters, our simulation reproduced the experimental data in other experiments used in this study (**Figure 4**, **S1D-F**). Although it is mechanistically not precise, our heuristic model could nevertheless provide insights into the nature of the replisome dynamics. Our simulations provided the following estimates (mean ± standard error on mean):

DNAP load time constant = 4.7 ± 0.4 min Low temperature DNAP rate = 27 ± 3 min High temperature rate = 57 ± 4 min DNAP_stall_time_constant = 64 ± 20 min Fraction pre-pre-existing DNAP = 20 ± 2 %

### Framework for simulation of SMC extrusion on replicated chromosomes

To assess the possible rules of interaction between loop-extruding SMC complexes and replisomes, we extended our previous framework for simulations SMC-SMC interactions ^14^ (https://github.com/hbbrandao/bacterialSMCtrajectories) by including replicated chromosomes and SMC-replisome interactions.

Each simulated chromosome was segmented into 4040 bins of 1 kb each to resemble the total length ∼4033 kb in our bacterial strains and to best match the number of 404 bins of 10 kb used for Hi-C and other analyses. In addition to the 4040 bins for the main (unreplicated) chromosome present in all simulations, we also added a number of lattice sites for the segments of chromosome that have been replicated. As such, each simulation could contain up to 4040 ⨉ 2 = 8080 chromosomal lattice sites to allow for up to two whole replicated chromosomes per cell. At the beginning of each simulation, replisome positions were assigned based on *fraction_pre_existing_DNAP* (see above).

For all simulations, we allowed the number of SMCs on the chromosome to fluctuate based on their loading/unloading dynamics, which was done by including a “cytosolic lattice site” for SMC to reside in addition to the aforementioned chromosomal lattice sites. We used a constant number of 40 SMC loop-extruding molecules per cell as determined by previous experiments and simulations ^14^. At the start of each simulation, ∼75% of SMCs (30 loop extruders) started off at the “cytosolic” locus to account for the fact that SMCs have not yet been induced to load on the DNA. ∼25% (10 loop extruders) were “pre-loaded” throughout the chromosome to mimic the non-specific loading of SMCs in the absence of *parS* sites or ParB protein ^14, 15^.

For simulating SMC dynamics, we created a Python program defining a smcTranslocator class as the engine for simulating the motion of SMC complexes on the replicating chromosome. This class encapsulated the dynamic behaviors of SMCs, including their loading onto the DNA, their translocation along the chromosome, and their unloading. Broadly, the simulation of SMC dynamics can be broken down into three main parts ordered by when they occur in each simulation time step: **death**, **birth** and **step**: **death**: This function governs the unloading or dissociation of SMCs from the chromosome. It includes both the basal unloading rate which represents the intrinsic tendency of SMCs to dissociate from the DNA, and the possibility of unloading triggered by interactions with other SMC complexes or interactions with the replisome. Death probabilities were pre-set at the start of each simulation and were calibrated for each simulation time-step to specific average lifetimes for the SMCs at each type of lattice site. For the cytosolic lattice site, the average lifetime was calibrated to be 180 s. For the chromosomal lattice sites including the “*parS* lattice site” and “regular lattice sites”, the average lifetime of the SMC was set to 1200 s as determined from experiments (see the SMC lifetime calibration section below for details). For “replication fork lattice sites”, the average lifetime (or probability of unloading) was left as a free parameter for each simulation (i.e. we performed parameter sweeps over this value with values ranging from 1 s [the case of immediate unloading] to 10,000,000 s [the case of no unloading]). Finally, we also defined a set of 100 terminus lattice sites (i.e. lattice sites 1950 to 2050 at the replication terminus region) to mimic the unloading of SMCs ^16^. An SMC at a *ter* site had a probability of unloading of 10% per simulation time step.

**birth**: This function orchestrates the transfer of dissociated SMCs onto one of the three possible simulated lattice sites described above (i.e. two for the chromosome, or one for the cytoplasm). The loading probabilities are pre-set at the start of each simulation by defining lattice sites as one of: “cytosolic” (loading strength 16,000), “*parS*” (loading strength 4000), “replication fork” (loading strength 1) or “regular” (loading strength 1). As an example, for a fully replicated chromosome has 8080 chromosomal lattice sites, including two *parS* sites (one for each daughter DNA molecules), and 2 ⨉ 4039 regular chromosomal lattice sites, separate from the lattice site for cytoplasmic SMCs. In this example, an SMC undergoing the “birth” (or loading) operation would be randomly selected to bind to the cytoplasmic lattice site with probability 16000/(16000 + 2 ⨉ 4000 + 4039 ⨉ 1 ⨉ 2) = 49.9%, to a *parS* site with probability (2 ⨉ 4000)/(16000 + 2 ⨉ 4000 + 4039 ⨉ 1 ⨉ 2) = 24.9%, and to any other regular site with probability 1/(16000 + 2 ⨉ 4000 + 4039 ⨉ 1 ⨉ 2) = 0.003% .

**step**: This function simulates the translocation of SMCs along the chromosome. For each simulation step, it considers various factors influencing SMC movement, such as directionality, translocation rate and stalling probabilities. The step function uses periodic boundary conditions to model SMC movement on a circular chromosome. When an SMC moves past one end, it is repositioned to the other, creating a continuous track for translocation, reflecting the circular nature of bacterial chromosomes. In our simulations, SMC loop extruders consist of two “motor subunits” that can move independently from one another ^17^, and interact independently with other SMCs and the replisome. By default, the interaction rule of an SMC with a replisome is “pausing” (also known as blocking or doing nothing). If an SMC subunit translocating on unreplicated DNA in *ter*-to-*ori* orientation needs to bypass (step over) a lattice site containing the replication fork (head-on collisions between SMC and replisome), the “step” function randomly selects one of the two newly replicated daughter DNA strands to move to with probability of 50% for either one. For the reverse cases where an SMC subunit moving on newly replicated DNA strands in an *ori-to-ter* direction encounters a replication fork site ahead (head-to-tail collision), we assume SMCs move from the daughter DNA to the unreplicated mother DNA (i.e. transitions from daughter DNA to daughter DNA are not allowed). For the rules of engagement of SMCs with other SMCs, we fixed the rates of pausing (probability of 0.9275 per simulation step), bypassing (probability of 0.07 per simulation step) and facilitated dissociation (probability of 0.0025 per simulation step) as previously described ^14^. Lastly, to best mimic the asymmetric translocation dynamics of the SMC complexes, we specified a genomic bias for the translocation: SMCs translocating from lattice site 90 (i.e. position 90 kb from ori) towards the terminus (lattice site 2000) move at the “full speed”, and from lattice site 2000 to 90 at 33% of the full speed to reproduce the correct “tilt” in the Hi-C secondary diagonal ^17^. The full speed of SMC used in the simulations was 1.07 kb/s at 42°C, close to the value of 1.18 kb/s as measured by experimental Hi-C (**Figure S3B**), and was 0.35 kb/s at 30°C as inferred by fitting a time-course of SMC ChIP-seq data (**Figure S2**). All simulation time step units are calibrated to an equivalent of one second of real experimental time.

We note that all of the above simulations of SMC extrusion dynamics are one dimensional in nature.

### Calibration of spontaneous dissociation rate of SMC complexes

We constrained the spontaneous dissociation rate of SMCs at 42°C using SMC ChIP-seq (**Figure S4**). We used a strain (BWX4504) containing a single *parS* site at -1° and IPTG-inducible *sirA* which arrested cells at G1 after one-hour treatment with 1 mM IPTG at 42°. In this dataset, most of the chromosomes were unreplicated, but the MFA analysis suggested that a small fraction of chromosomes in the cell population had pre-existing replisomes in a similar way as seen in MFA data of the *dnaB (ts)* strain described above (see the section *calibration of replisome dynamics model)* (**Figure S1A**, slope 1). As such, we took several steps to ensure that our estimate of the spontaneous SMC dissociation rate was not strongly biased by this fraction of pre-existing replisomes. We took the following key steps to determine the spontaneous SMC dissociation rate:

1. **Data preparation:** Loading and binning experimental ChIP-seq and MFA data.
2. **Replisome simulation:** Because the cells were arrested in G1, we set the DNAP loading time to 100000 min, which was effectively not allowing any replisomes to load on the DNA. Using the fitting procedure we described in Calibration of replisome dynamics model, we identified that 5% of chromosomes had pre-existing replisomes. We considered their effects on SMC dynamics (see below).
3. **SMC translocation simulations:** We implemented a function (smcTranslocator) to simulate SMC movement on the chromosome, incorporating factors like bypassing, unloading, and pausing at the replication fork positions. We performed four simulations to account for the 5% of pre-existing replisomes. We first simulated the three extreme interaction scenarios of between SMC and the replisome: 1) “blocking” where SMC translocation is blocked by the replisome and SMC remains at the collision site; 2) “unloading” where SMC immediately dissociates from the chromosome when encountering the replisome; 3) “bypassing” where SMC immediately bypasses the replisome. We reasoned that if all the fit values for these extreme scenarios gave similar “best-fits”, then the residual 5% of pre-existing replisomes would have an overall small effect on the estimated SMC dissociation rate. As a final sanity check, we ran a fourth set of simulation assuming there was no residual (i.e. partially replicated) chromosome.
4. **Parameter sweep:** We systematically varied the SMC lifetime (i.e. 1/ (spontaneous dissociation rate)) to generate a series of simulated SMC enrichment profiles. We swept lifetime values from 200 s to 2400 s at intervals of 200 s. The resulting simulated SMC profiles were then compared directly to the ChIP-seq experimental data (**Figure S4A**). We inspected the results both visually and by computing a goodness of fit metric.
5. **Goodness-of-fit analysis:** We compared simulated SMC enrichment profiles with experimental data to identify the SMC dissociation rate. The goodness-of-fit metrics we employed was the mean absolute deviation between the simulated SMC profiles and the ChIP-seq data (**Figure S4B**). By considering the goodness-of-fit curves for all three extreme-value scenarios of SMC-replisome interaction rules, we found that the overall best fit value for the average SMC lifetime was ∼1200 s.

### Generation of SMC occupancy profiles from simulations

Each SMC loop extruder has two motors (the “left” extrusion motor and the “right” extrusion motor). Thus, at each simulation time step, the position of one SMC loop extruder has a “pair” of positions corresponding to the two motors. For each lattice site, we counted the number of instances when a SMC motor was on it. The counts were summed for all SMCs on all chromosomes in a simulation (typically at least 1025 chromosomes) to produce the plotted SMC occupancy profiles (i.e. simulated ChIP). As done for experimental ChIP-seq data, we discretized the simulated genome into 404 bins (each consisting of 10 lattice sites) and plotted the median SMC occupancy count value within each 10 kb bin. Lastly, we normalized the entire distribution by the area under the curve (i.e. the sum of all 404 median bin counts) and multiplied the result by 4200 to report the simulated SMC occupancy profile in reads per million. In this way, we can directly compare the simulated SMC enrichments to the experimental ChIP enrichments.

### Generation of Hi-C maps from simulations

For generating Hi-C maps from simulations, we used the semi-analytical methods described in Brandao et al, 2021 ^14^ which builds on the work from Banigan et al, 2020 ^18^. Briefly, the simulated Hi-C maps come from calculations of the contact probabilities between different parts of the chromosome (i.e. the chromosomal lattice sites). Sine these contacts are mediated by the two motors of SMCs, the method used positions of the loop extruding SMC complexes on the genome to compute an undirected graph to obtain the shortest path between any two randomly sampled chromosomal lattice sites. We used Scipy1.4.1 (scipy.sparse.csgraph.shortest_path) for the shortest path calculation.

To create the simulated Hi-C maps shown in the **Figure 5**, we sampled at least 5152 chromosomes for each map, with 1000000 randomly sampled unique contact pairs for each chromosome. Sections of the genome with replicated chromosomes were averaged. Contacts were binned to 10 kb resolution, resulting in a single Hi-C contact map of 404 bins, which was comparable to the experiments. We note that it was not possible to directly compare the Hi-C contact values in the simulated map with those in the experimental data because of the shortest-path approximation ^14^, Gaussian contact model approximation ^18^, and the lack of simulated “fine structures” such as plectonemes ^19^. To closely match the experimental contact probability decay and to show the Hi-C maps on comparable color scales, we adjusted the scaling exponent in the Hi-C simulations. This adjustment aids visual comparison but does not alter our results or conclusions.

### Filtering of ChIP-seq and MFA data for goodness-of-fit calculations and visual comparison

All ChIP-seq and MFA data were mapped to 1 bp resolution. We then applied a 10 kb moving “window” and measured the median value within the window. Data were discretized to 404 bins to cover the entirety of the genome similarly to the process described above for simulated SMC occupancy profiles. For direct comparisons of the simulated SMC enrichment profiles to experimental ChIP-seq data, we did not normalize the ChIP by the DNA input (i.e. MFA data) because of two reasons: 1) our simulations already account for the distribution of multiple chromosome copies; 2) since MFA data are noisy, we reasoned that comparing the ChIP-seq data directly to the simulated profiles would result in higher confidence and less bias.

### Parameter sweeps for interactions rules between SMC and replisome

We took the following steps to generate parameter sweeps:

1. **Data preparation:** We loaded and binned experimental ChIP-seq and MFA data as described above. We fit the data using one timepoint specified in the figure legend (e.g. we used +IPTG 25min in **Figure 5A** and **F**).
2. **Replisome simulation:** To mimic the distribution of replisomes for each experimental condition, we used the best-fit values that reproduced the MFA distributions of cells in Figure 1C (see the section *Calibration of replisome dynamics model*, also see **Figure S1BC**), which also reproduced the MFA plots for the stalled replisomes (**Figure S1D-F**).
3. **SMC translocation simulations and parameter sweep:** We employed the smcTranslocator function to simulate SMC movement on the chromosome, incorporating factors like bypassing, unloading, and blocking (i.e. pausing) at the replication fork positions. While keeping all other parameters fixed (as described above in the section *Framework for simulation of SMC extrusion on replicated chromosomes*), we systematically varied the *SMC_bypassing_time* and the *SMC_unloading_time* values. We used all pairwise combinations of the following times [3.75, 7.5, 15, 30, 60, 120, 240, 480] in seconds. Additionally, to simulate the simple models, we used the pairs of values for [*SMC_bypassing_time; SMC_unloading_time*] of i) [10,000,000; 10,000,000] for blocking only; ii) [1; 10,000,000] for immediate bypassing only; and iii) [10,000,000; 1] for immediate unloading only. We median-filtered the resulting simulated SMC profiles as done for the experiments, and then compared the simulations and experiments directly both visually and by computing a goodness-of-fit metric (**Figures S5** and **S6**).
4. **Goodness-of-fit analysis:** We compared simulated SMC profiles with experimental data to identify the parameter combination that best explains the observed SMC distribution (**Figures S5B** and **S6B**). The goodness-of-fit metric we employed was the mean absolute deviation between the simulated SMC profiles and the experimental ChIP-seq data.

